# Contingency and chance erase necessity in the experimental evolution of ancestral proteins

**DOI:** 10.1101/2020.08.29.273581

**Authors:** Victoria Cochran Xie, Jinyue Pu, Brian P.H. Metzger, Joseph W. Thornton, Bryan C. Dickinson

## Abstract

To understand why evolution produced the biological systems that exist today, we must know how important chance, contingency, and necessity were during history. Previous observations suggest that each of these modes of causality affects evolution in various settings, but their relative roles and interactions are not well characterized because they have never been systematically assessed in a single system or on a timescale relevant to evolutionary history. To this end, we reconstructed ancestral B-cell-lymphoma-2-family proteins and developed a continuous evolution method to select for defined protein-protein interaction specificities. By repeatedly evolving a series of ancestral proteins to acquire specificities that occurred during history, we show that contingency steadily overwhelms chance and erases necessity as the primary cause of sequence variation in proteins over long phylogenetic timescales. As a result, evolutionary trajectories launched from distant starting points are essentially unpredictable, even under strong and identical selection pressures. Genetic dissection of the outcomes shows that chance arises because numerous sets of mutations can alter specificity at any point in time, while contingency arises because historical substitutions change these sets. Patterns of variation in extant protein sequences are therefore largely the idiosyncratic product of a particular course of unpredictable historical events.

## INTRODUCTION

The extent to which biological diversity is the necessary result of optimization by natural selection or the unpredictable product of random events and historical contingency is one of evolutionary biology’s most fundamental and unresolved questions (Gould 1990; Travisano et al. 1995; Ramsey and Pence 2016; Jablonski 2017). The answer to this question would have strong implications not only for our understanding of evolutionary processes but also for how we should analyze the particular forms of variation that exist in nature today. For example, if diversity primarily reflects a predictable process of adaptation to distinct environments, then a central goal of biology would be to explain how the characteristics of living things help to execute particular functions and improve fitness (Mayr 1983). By contrast, if diversity reflects chance sampling from a set of similarly fit possibilities, then the variation itself is of little interest because it does not affect biological properties or shape future evolutionary outcomes; the goal of biology would be to identify invariant characteristics of natural systems and explain how they contribute to function (Monod 1972; Kimura 1983; Lobkovsky and Koonin 2012). Finally, if diversity reflects contingency—a strong dependence of future events on past and current states—then the outcomes of evolution would be predictable only given complete knowledge of the constraints and opportunities specific to each starting point (Gould and Lewontin 1979; Beatty 2009; Blount et al. 2018); the goal of biology would then be to characterize these constraints and opportunities, their mechanistic causes, and the historical events that shaped them.

Many studies have provided insight into the ways that chance, contingency, and necessity can affect evolution, but the relative importance of these factors during evolutionary history remains unresolved because they have never been measured in the same system, and their effects over long evolutionary time scales have not been characterized. For example, experiments on ancestral proteins have shown that particular historical mutations have different effects when introduced into different ancestral backgrounds – suggesting contingency – but they do not reveal the extent to which context-dependence actually influenced evolutionary outcomes; further, these historical trajectories happened only once, so they cannot elucidate the effect of contingency relative to chance (Ortlund et al. 2007; Bridgham et al. 2009; Bloom et al. 2010; Gong et al. 2013; Harms and Thornton 2014; McKeown et al. 2014; Risso et al. 2015; Natarajan et al. 2016; Starr et al. 2018; Wu et al. 2018). Experimental evolution studies could, in principle, characterize both chance and contingency if they had sufficient replication from multiple starting points, but to date no study has done so; further, historically relevant proteins have not been used as starting points and historically relevant functions have not been selected for, so these studies’ relevance to historical evolution is not clear (Wichman et al. 1999; Couñago et al. 2006; Bollback and Huelsenbeck 2009; Salverda et al. 2011; Blount et al. 2012; Meyer et al. 2012; van Ditmarsch et al. 2013; Dickinson et al. 2013; Spor et al. 2013; Kryazhimskiy et al. 2014; Wünsche et al. 2017; Kacar et al. 2017; Baier et al. 2019; Zheng et al. 2019). Studies of phenotypic convergence in nature suggest some degree of repeatability at the genetic level (reviewed in (Arendt and Reznick 2008; Gompel and Prud’homme 2009; Orgogozo 2015; Storz 2016)), but these studies rarely involve replicate lineages from the same starting genotypes, and evolutionary conditions are seldom identical; similarities and differences among lineages can therefore not be attributed to chance, contingency, or necessity. Further, these studies have typically involved closely related species or populations and therefore do not measure the effects of chance and contingency that might be generated during long-term evolution.

The ideal experiment to determine the relative roles of chance, contingency, and necessity in historical evolution would be to travel back in time, re-launch evolution multiple times from each of various starting points that existed during history, and allow these trajectories to play out under historical environmental conditions (Gould 1990). By comparing outcomes among replicates launched from the same starting point, we could estimate the effects of chance; by comparing those from different starting points, we could reveal the effects of contingency. Necessity would be apparent if both chance and contingency were absent, with the same outcome occurring repeatedly, irrespective of the point in history from which evolution was launched (Figure 1A). Although time travel is currently impossible, we can come close to this ideal experimental design by reconstructing ancestral proteins as they existed in the deep past (Thornton 2004) and using them to launch replicated evolutionary trajectories in the laboratory under selection to acquire the same molecular functions that evolved during history.

**Figure 1.**
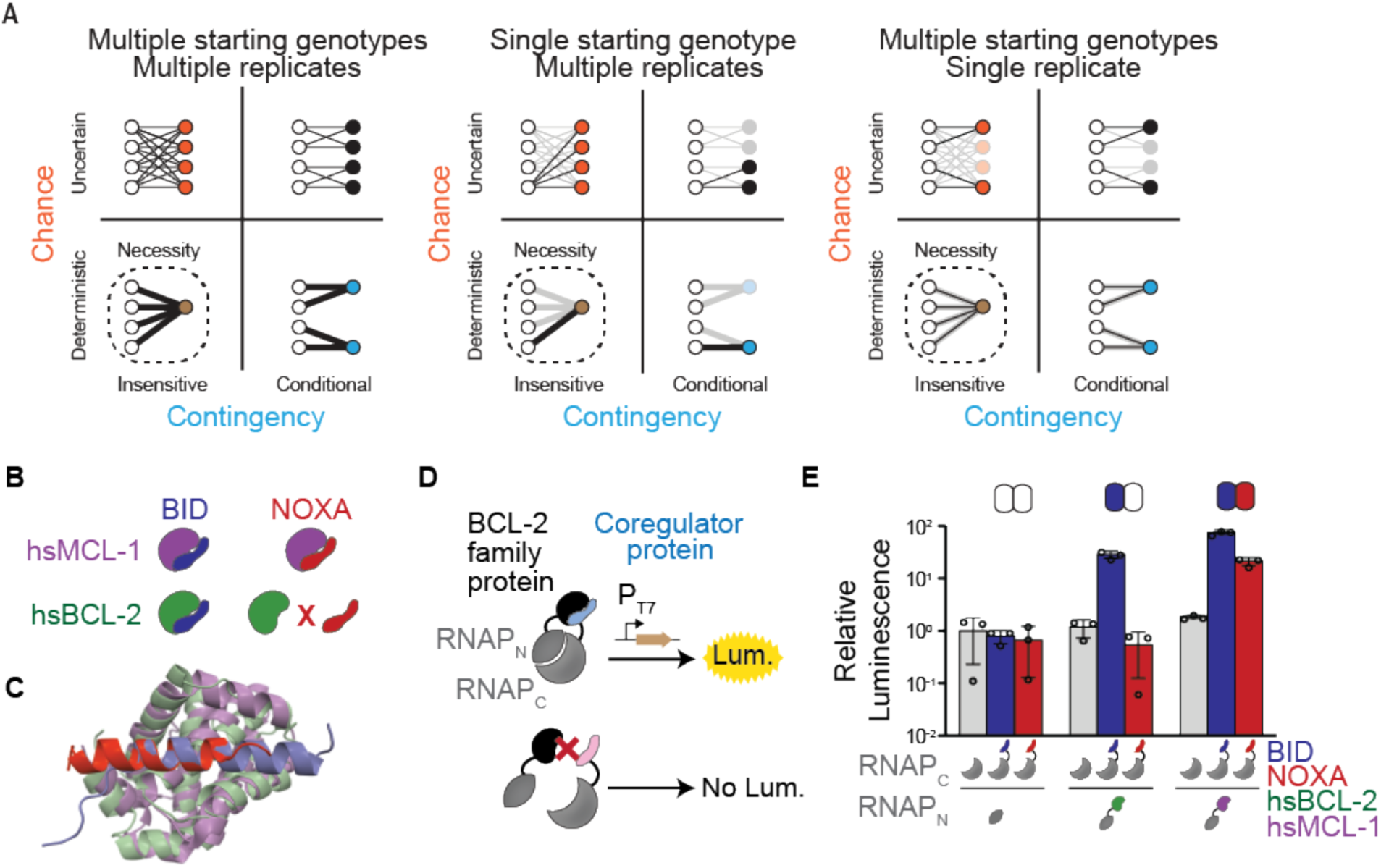
Assessing chance and contingency in the BCL-2 family of proteins. (A) The outcomes of evolution can be influenced by chance (y-axis) and/or contingency (x-axis). Black lines connect starting genotypes (white circles) to evolutionary outcomes; thickness is proportional to a path’s probability. Each cluster depicts evolutionary outcomes under the influence of chance (orange), contingency (blue), or both (black); outcomes are necessary (brown, with dotted line) when neither is important. *Left:* evolution experiments in which replicate trajectories are initiated from multiple starting points can distinguish chance and contingency. *Middle*, multiple replicates from one starting point can detect chance but not contingency, because outcomes from other starting points are not observed (faded lines and circles). *Right*, initiating one replicate from multiple starting points underestimates chance. (B) Protein binding specificities of extant BCL-2 family members. Human MCL-1 (hsMCL-1, purple) binds BID (blue) and NOXA (red), while human BCL-2 (hsBCL-2, green) binds BID but not NOXA. (C) Crystal structures of MCL-1 (purple) bound to NOXA (red, PDB 2nla), and BCL-xL (green, a closely related paralog of BCL-2) bound to BID (blue, PDB 4qve). (D) Schematic of the luciferase reporter assay to assess PPIs. If a BCL-2 family protein (black) binds a coregulator protein (blue), the split T7 RNAP biosensor (gray) assembles and drives luciferase expression. If a coregulator (pink) is not bound, no luciferase is expressed. (E) Interactions of human BCL-2 and MCL-1 with BID (blue bars) and NOXA (red) in the luciferase assay, compared to no-coregulator control (gray). Activity is scaled relative to no-coregulator control with no-BCL-2 protein. Columns and error bars, mean ± SD of three biological replicates (circles). Shaded boxes above show the same data in heatmap form: BID activity is normalized relative to hsBCL-2 with BID; NOXA activity is normalized to hsMCL-1 with NOXA.

Here we implement this strategy using the B-cell lymphoma-2 (BCL-2) protein family as a model system and PPI specificity as the target of natural selection. BCL-2 family proteins are involved in the regulation of apoptosis (Danial and Korsmeyer 2004; Petros et al. 2004; Chipuk et al. 2010; Kale et al. 2018) through specific PPIs with coregulators (Chen et al. 2005; Lomonosova and Chinnadurai 2008; Dutta et al. 2010). Among BCL-2 family members, the Myeloid Cell Leukemia Sequence 1 (MCL-1) class binds both the BID and NOXA coregulators, whereas the BCL-2 class (a subset of the larger BCL-2 protein family) binds BID but not NOXA (Figure 1B) (Certo et al. 2006). The two classes share an ancient evolutionary origin: both are found throughout the Metazoa (Lanave et al. 2004; Banjara et al. 2020) and are structurally similar, using the same cleft to interact with their coregulators (Figures 1C and S1), despite having only 20% sequence identity.

To drive the evolution of new PPI specificities, we developed a novel high-throughput Phage Assisted Continuous Evolution (PACE) system (Esvelt et al. 2011) for simultaneous selection for and against particular PPIs (Pu et al. 2017b, 2019). We applied this technique to a series of reconstructed ancestral BCL-2 family members, repeatedly evolving each starting genotype to acquire PPI specificities found among extant family members. By analyzing sequence outcomes among replicates from the same and different starting points, we were able to quantify the roles of chance, contingency, and necessity in the evolution of PPI specificity, characterize how they have changed over time, and dissect the underlying genetic basis by which these properties emerge.

## RESULTS

### BID specificity is derived from an ancestor that bound both BID and NOXA

We first characterized the historical evolution of PPI specificity in the BCL-2 family using ancestral protein reconstruction. We inferred the maximum likelihood phylogeny of the family, recovering the expected sister relationship between the metazoan BCL-2 and MCL-1 classes (Figure 2, Figure S2A). We then reconstructed the most recent common ancestor (AncMB1) of the two classes, which represents a gene duplication that occurred before the last common ancestor of all animals, and 11 other ancestral proteins that existed along the lineages leading from AncMB1 to human BCL-2 (hsBCL-2) and to human MCL-1 (hsMCL-1) (Supplementary Table S1).

**Figure 2.**
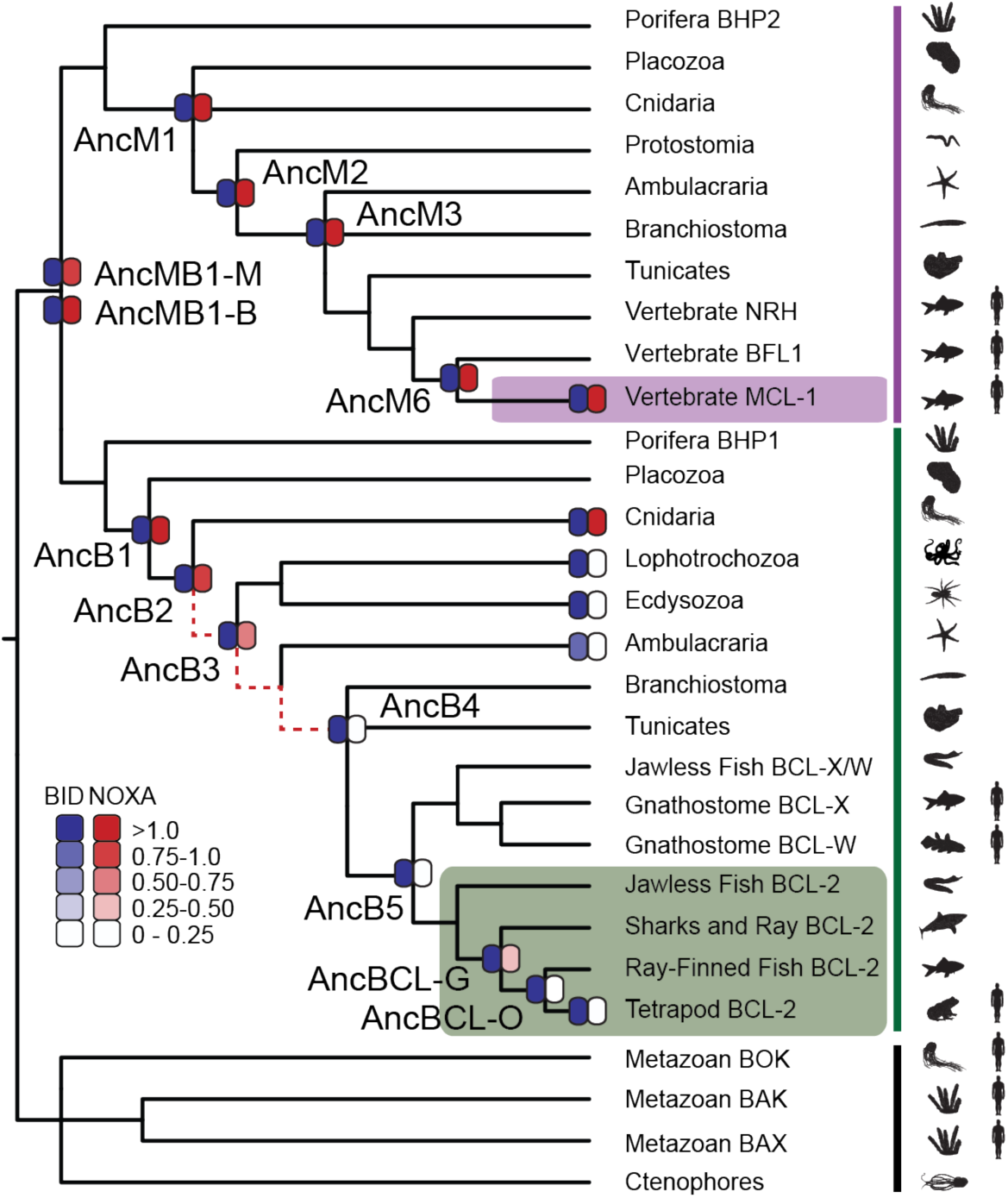
BID specificity was acquired during vertebrate BCL-2 evolution. Reduced maximum likelihood phylogeny of BCL-2 family proteins. Purple bar, MCL-1 class; green bar, BCL-2 class. The phylogeny was rooted on outgroup using paralogs BOX, BAK, and BAX (black bar). Heatmaps indicate BID (blue) and NOXA (red) binding measured using the luciferase assay as in Figure 1. Each shaded box shows the normalized mean of three biological replicates. Red dotted lines, interval during which NOXA binding was lost, yielding BID specificity in the BCL-2 protein of vertebrates (green box). Purple box, vertebrate MCL-1. Silhouettes, representative species in each terminus. AncMB1-M and -B are alternative reconstructions using different approaches to alignment ambiguity (see Methods).

We synthesized genes coding for these proteins and experimentally assayed their ability to bind BID and NOXA using a proximity-dependent split RNA polymerase (RNAP) luciferase assay (Figures 1D and 1E) (Pu et al. 2017b). AncMB1 bound both BID and NOXA, as did all ancestral proteins in the MCL-1 clade and hsMCL-1 (Figure 2, Figure S2, Table S1). Ancestral proteins in the BCL-2 clade that existed before the last common ancestor of deuterostomes also bound both BID and NOXA, whereas BCL-2 ancestors within the deuterostomes bound only BID, just as hsBCL-2 does. This reconstruction of history was robust to uncertainty in the ancestral sequences: experiments on “AltAll” proteins at each ancestral node – which combine all plausible alternative amino acid states (PP>0.2) in a single “worst-case” alternative reconstruction – also showed that BID specificity arose within the BCL-2 clade (Table S2).

To further test this inferred history, we characterized the coregulator specificity of extant BCL-2 class proteins from taxonomic group in particularly informative phylogenetic positions. Those from cnidaria were activated by both BID and NOXA, whereas those from protostomes and invertebrate deuterostomes were BID-specific (Figure 2, Figure S2, Table S1). These results corroborate the inferences made from ancestral proteins and indicate that BID specificity evolved when the ancestral ability to bind NOXA was lost between AncB2 (in the ancestral eumetazoan) and AncB4 (in the ancestral deuterostome).

### A directed continuous evolution system for rapid changes in PPI specificity

To rapidly evolve BCL-2 family proteins to acquire the same PPI specificities that existed during the family’s history, we developed a novel phage-assisted continuous evolution (PACE) system (Esvelt et al. 2011) (Figures 3A, S3A, and S3B). Previous PACE systems have evolved binding to new protein partners using a bacterial 2-hybrid approach (Badran et al. 2016), but evolving PPI specificity per se requires simultaneous selection for a desired PPI and against an undesired PPI. For this purpose, we used two orthogonal proximity-dependent split RNAPs that recognize different promoters in the same cell. The N-terminal fragment of RNAP was fused to the BCL-2 protein of interest and encoded in the phage genome, and two C-terminal RNAP fragments (C-RNAP), each fused to a different BCL-2 co-regulator, were encoded on host plasmids: one C-RNAP, fused to the selected-for coregulator, drives expression of an essential viral gene (gIII) when reconstituted by binding to the BCL-2 protein, while the other C-RNAP, fused to the counter-selected coregulator, drives expression of a dominant-negative version of gIII (Pu et al. 2017a). After optimizing this system, we used activity-dependent plaque assays and phage growth assays to confirm that it can impose strong selection for the PPI specificity profiles of extant hsBCL-2 and hsMCL1 (Figures 3B and 3C).

**Figure 3.**
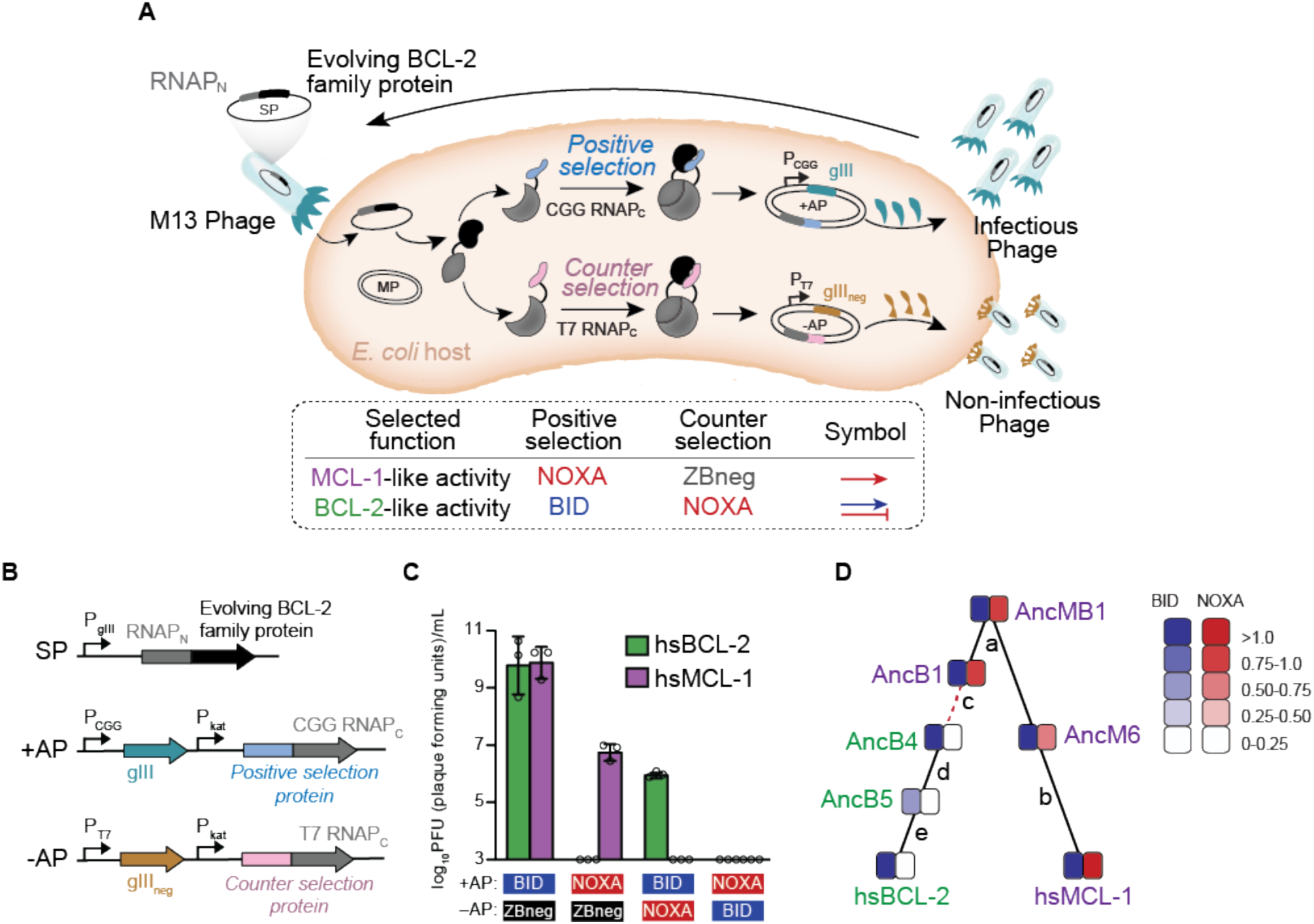
Continuous directed evolution of modern and ancestral BCL-2 family proteins. (A) PACE system for evolving specificity in PPIs. A BCL-2 family protein (black) is fused to the N-terminal portion of T7 RNA polymerase (RNAP_N_, gray) and placed in the phage genome (SP). Host *E. coli* carry a positive selection accessory plasmid (+AP), containing gIII (teal) driven by a T7_CGG_-promoter (P_CGG_) and the C-terminal portion of T7_CGG_ RNAP (CGG RNAP_C_) fused to the target binding protein (light blue). Target binding reconstitutes RNAP_CGG_ and drives expression of gIII, generating infectious phage. The counterselection accessory plasmid (-AP) contains dominant negative gIII (gIII_neg_, gold) driven by a T7 promoter (P_T7_) and the C-terminal portion of T7 RNA polymerase (T7 RNAP) fused to a counterselection protein. An arabinose-inducible mutagenesis plasmid (MP) increases the mutation rate. To evolve BCL-2 like specificity, positive selection to bind BID was imposed with counterselection to avoid binding NOXA (blue arrow and red bar). To evolve MCL-1 like activity, positive selection to bind NOXA (red arrow) was imposed with counterselection to avoid nonspecific binding using a control zipper peptide (ZBneg). (B) Map of the phage selection plasmid (SP) and the positive and counterselection accessory plasmids (+AP and -AP). hsBCL-2 (green). hsMCL-1 (purple). (C) Growth assays to assess selection and counterselection. PFU is shown after culturing 1000 phage containing hsBCL-2 (green) or hsMCL-1 (purple) on *E. coli* containing various APs. Detection limit 10^3^ PFU/mL. Bars show mean ± SD of three replicates (circles). (D) Phylogenetic relationship among starting genotypes for experimental evolution. Each starting protein was evolved in multiple independent replicate for new PPI specificity. Green, proteins selected to gain NOXA binding; purple, proteins selected to lose NOXA binding. Heatmaps, luciferase assay activity (mean of three biological replicates) of starting proteins. Red dashed line, interval during which NOXA binding was historically lost, yielding BID specificity in the BCL-2 clade. Letters, index of phylogenetic intervals between ancestral proteins referred to in Fig. 5.

The simplicity of this platform allowed us to drive extant and reconstructed ancestral proteins to recapitulate or reverse the historical evolution of the BCL-2’s binding PPI specificity in multiple replicates in just days, without severe experimental bottlenecks (Figure 3D). Three proteins with the ancestral phenotype—hsMCL-1, AncM6, and AncB1—were selected to acquire the derived BCL-2 phenotype, losing NOXA binding but retaining BID binding (Figure S3C). Conversely, hsBCL-2, AncB5, and AncB4 were evolved to revert to the ancestral phenotype, by gaining NOXA binding (Figure S3D). For each starting genotype, we performed four replicate experimental evolution trajectories (Table S3). Each experiment was run for four days, corresponding to approximately 100 rounds of viral replication (Esvelt et al. 2011). All trajectories yielded the target PPI specificity, which we confirmed by experimental analysis of randomly isolated phage clones using activity-dependent plaque assays and *in vivo* and *in vitro* binding assays (Figures 4A-B and S3E-L).

**Figure 4.**
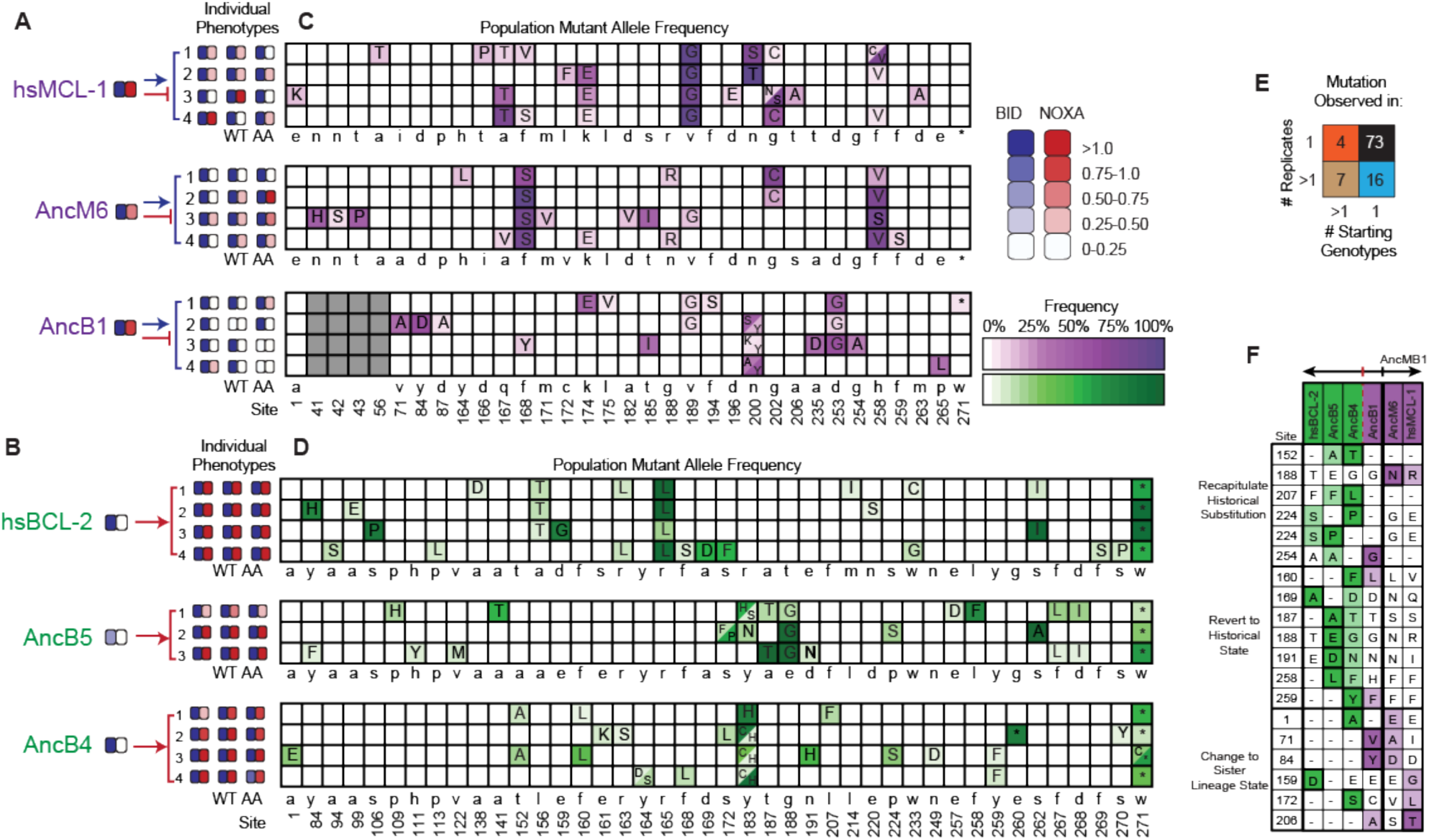
Chance and contingency shape evolutionary outcomes. (A) Phenotypic outcome of PACE experiments when proteins with MCL-1-like specificity were selected to maintain BID and lose NOXA binding. For each starting genotype, the BID (blue) and NOXA (red) binding activity of the starting genotype and three phage variants isolated from each evolved replicate (number) are shown as heatmaps. (B) Phenotypic outcome of PACE experiments when proteins with BCL-2-like specificity were selected to gain NOXA binding. (C) Frequency of acquired states in PACE experiments when proteins with MCL-1-like specificity were selected to maintain BID and lose NOXA binding. Rows, outcomes of each replicate trajectory. Columns, sites that acquired one or more non-wildtype amino acids (letters in cells) at frequency >5%; color saturation shows the frequency of the acquired state. Site numbers and wildtype amino acid (WT AA) states are listed. Gray, sites that do not exist in AncB1. (D) Frequency of acquired states when BCL-2-like proteins were selected to gain NOXA binding. (E) Repeatability of acquired states across replicates. The 100 non-WT states acquired in all experiments were categorized as occurring in 1 or >1 replicate trajectory from 1 or >1 unique starting genotype, with the number in each category shown. The vast majority of states evolved in just one replicate from one starting point (black). (F) States acquired during PACE that also occurred during evolutionary history. Rows, sites at which a state was acquired at >5% frequency in a PACE experiment that is also found in one or more of the ancestral or extant proteins (columns). Dark shaded cells, wild-type state in the protein used as the starting point for the PACE trajectory. Light shaded cells, acquired PACE state in the column for the historical protein in which the state occurs. Unshaded cells with dashes have the same state as the PACE starting point. Purple, proteins with PPI specificity like MCL-1; green, like BCL-2. PACE mutations are grouped into those that i) recapitulate historical substitutions by acquiring a derived state that evolved in history in a descendant of the ancestor used as the PACE starting point; ii) reverse historical substitutions by acquiring a state from an ancestor earlier than the PACE starting point; or iii) confer a state acquired in history along a different lineage.

### Chance and contingency erase necessity in the evolution of PPI specificity

We used deep sequencing to compare the sequence outcomes of evolution across trajectories initiated from the same and different starting points. Necessity was almost entirely absent. Across all trajectories, 100 mutant amino acid states at 75 different sites evolved to frequency >5% in at least one replicate (Figures 4C-D and S4A-D and Table S4). Of these acquired states, 73 were states that appeared in only a single trajectory, and there were only 4 that arose in more than one replicate from multiple starting points (Figure 4E). No mutations were predictably acquired in all trajectories from all starting points when selection was imposed for binding to both BID and NOXA. The only mutation universally acquired under any selection regime was a nonsense mutation at codon 271, which was acquired in all trajectories selected for BID specificity, but experimental analysis of this mutation shows that it has no detectable effect on coregulator binding (Figure S4E).

Both chance and contingency contributed to this pervasive unpredictability. Pairs of trajectories launched from the same starting point differed, on average, at 78% of their acquired states, indicating a strong role for chance. Pairs that were launched from different starting points (but selected for the same PPI specificity) differed at 92% of acquired states, indicating an additional role for contingency. These starting points are separated by different amounts of evolutionary divergence, so to understand the extent of contingency on the timescale of metazoan evolution, we compared trajectories launched from AncB1 to those launched from hsMCL1 (the two most distant genotypes that were selected for BID specificity): of 34 states acquired in these experiments, only three occurred in at least one trajectory from both starting points. Of 40 states acquired in trajectories launched from AncB4 and hsBCL-2 (the two most distant proteins that were selected to gain NOXA binding), only one occurred in any trajectories from both starting points. Thus, contingency and chance together make sequence evolution in the BCL-2 family almost entirely unpredictable across long evolutionary timescales.

Consistent with a lack of necessity in the evolution of BCL-2 PPI specificity, virtually none of the mutations acquired in our experiments recapitulated or reversed substitutions that occurred during the historical interval when PPI specificity changed (Figures 4F and S5E-F). In all PACE trajectories that recapitulated the historical loss of NOXA binding, none acquired substitutions that occurred on the branch when NOXA binding was lost. Among all trajectories that reversed the historical loss of NOXA binding, only two ancestral states that were historically lost on the function-changing branch were acquired, and these both occurred in trajectories initiated from AncB4, the immediate daughter node of this branch.

### Contingency is the major determinant of sequence variation on long timescales

We next sought to directly quantify the relative effects of chance and contingency on the genetic outcomes of evolution and their relationship to phylogenetic divergence. We analyzed the genetic variance -- the probability that two variable sites, chosen at random, are different in state -- within and between trajectories from the same and different starting genotypes. To estimate the effects of chance, we compared the genetic variance between replicates initiated from the same starting genotype (V_g_) to the within-replicate genetic variance (V_r_). We found that V_g_ was on average 30 percent greater than V_r_, indicating that chance causes evolution to produce divergent genetic outcomes between independent lineages (Figure 5A). We quantified contingency by comparing the pooled genetic variance among replicates from different starting genotypes (V_t_) to that among replicates from the same starting genotype (V_g_). Contingency’s effect was even larger than that of chance, increasing V_t_ by an average of 80 percent across all pairs of starting points compared to V_g_. Together, chance and contingency had a multiplicative effect, increasing the genetic variance among trajectories from different starting genotypes (V_t_) by an average of 2.4-fold compared to the genetic variance within trajectories (V_r_).

**Figure 5.**
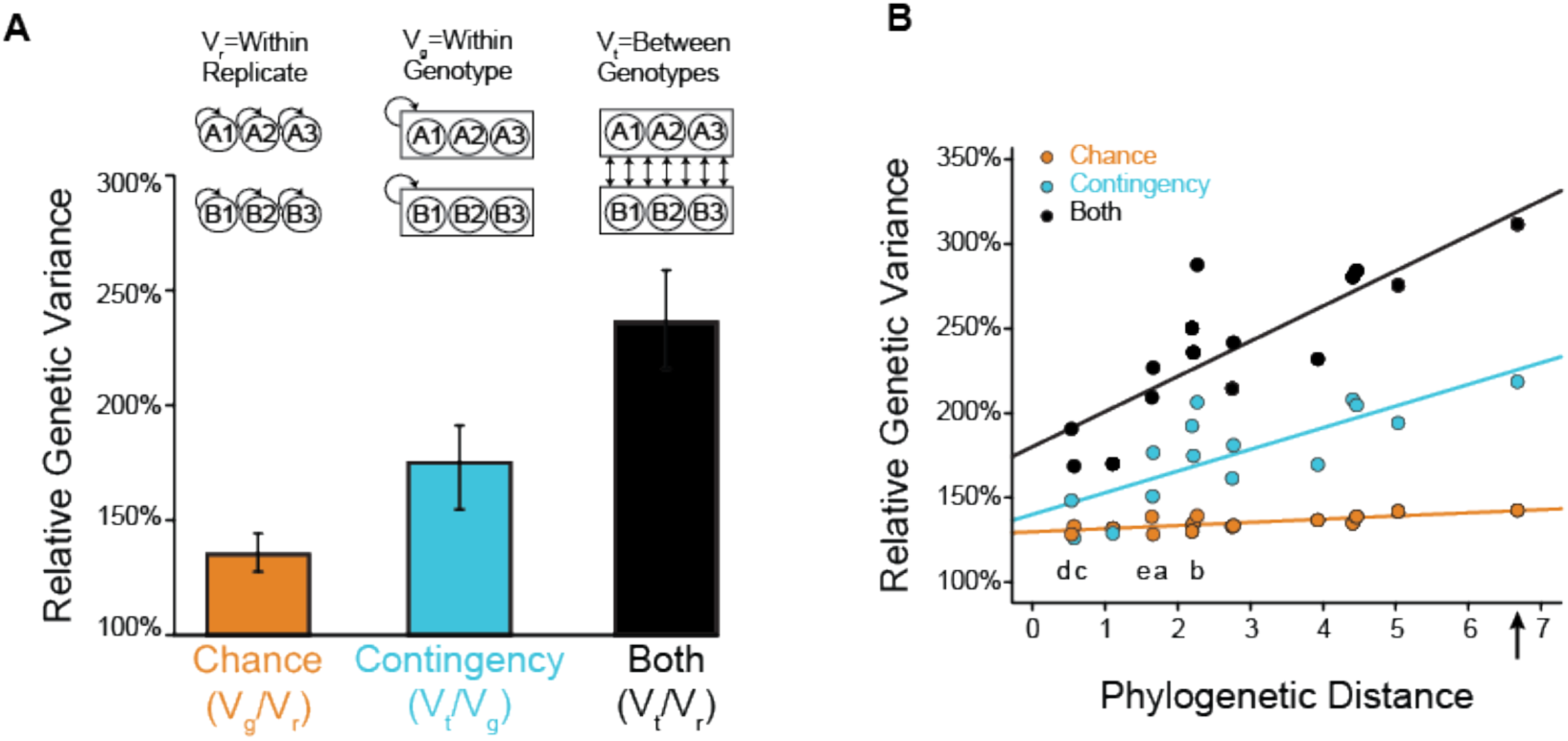
Effect of chance and contingency. (A) Variation in evolutionary sequence outcomes caused by chance (orange), contingency (teal), and both (black). Inset: schematic for estimating the effects of chance and contingency. Chance was estimated as the average genetic variance among replicates from the same starting genotype (V_g_) divided by the within-replicate genetic variance (V_r_). Contingency was estimated as the average genetic variance among replicates from different starting genotypes (V_t_) divided by the average genetic variance among replicates from the same starting genotype (V_g_). Combined effects of chance and contingency were estimated as the average genetic variance among replicates from different starting genotypes (V_t_) compared to the within replicate genetic variance (V_r_). Variance is the probability that two randomly drawn alleles are different in state. Error bars, 95% confidence intervals on the mean as estimated by bootstrapping PACE replicates. (B) Change in the effects of chance and contingency over phylogenetic distance. Each point is for a pair of starting proteins used for PACE, comparing the phylogenetic distance (the total length of branches separating them, in substitutions per site) to the effects of chance (orange), contingency (teal), or both (black), when PACE outcomes are compared between them. Solid lines, best-fit linear regression. Letters indicate the phylogenetic branch indexed in Figure 3D. The combined effect of chance and contingency increased significantly with phylogenetic distance (slope=0.19, p=2×10^−5^), as did the effect of contingency alone (slope=0.11, p=0.007). The effect of chance alone did not depend on phylogenetic distance (slope=0.02, p=0.5). The combined effect of chance and contingency increased significantly faster than the effect of contingency alone (0.08, p=0.04). Arrow, phylogenetic distance between extant hsMCL-1 and hsBCL-2 proteins, which share AncMB1 as their most recent common ancestor.

These comparisons do not account for phylogenetic structure or the extent of divergence between starting points. We therefore assessed how chance and contingency changed with phylogenetic distance using linear regression (Figures 5B and S5A-D). We found that the effect of contingency on genetic variance increased significantly with phylogenetic divergence among starting points. The effect of chance did not increase with divergence, but the combined effect of contingency and chance increased even more rapidly than contingency alone, because the total impact on genetic variance of these two factors is multiplicative by definition.

We next compared the impact of contingency to that of chance as phylogenetic divergence increases. On the timescale of metazoan evolution, contingency’s effect was three times greater than that of chance when evolution was launched from extant starting points that share as their last common ancestor AncMB1, near the base of Metazoa (Figure 5B). The combined effect of chance and contingency on this timescale was a 3.2-fold increase in variance among single trajectories launched from these starting points. Even across the shortest phylogenetic intervals we studied, contingency’s effect was larger than that of chance, although to a smaller extent. Taken together, these data indicate that contingency, magnified by the effect of chance, steadily increases the unpredictability of evolutionary outcomes as protein sequences diverge across history.

### Contingency is caused by epistasis between historical substitutions and specificity-changing mutations

Contingency in our system is expected to arise if historical substitutions (which separate ancestral starting points) interact epistatically with mutations that occur during selection-driven experimental evolution; this could cause the mutations that confer selected PPI specificities to differ among the starting points used for PACE. To experimentally test this hypothesis and characterize underlying epistatic interactions, we first identified sets of candidate causal mutations that arose repeatedly during PACE replicates from each starting genotype. We then verified their causal effect on specificity by introducing only these mutations into the protein that served as the starting point for the PACE experiment in which they were observed, and then measuring their effects on BID and NOXA binding: all were sufficient to confer the selected-for specificity in their “native” background.

We then introduced these mutations into the other starting proteins that had been subject to the same selection regime and performed the same assay (Figures 6A and 6B). Eleven of 12 such “swaps” failed to confer the PPI specificity on other proteins that they conferred in their native backgrounds. These swaps compromised binding of BID, failed to confer the selected-for gain or loss of NOXA binding, or both. The only case in which mutations that conferred the target phenotype during directed evolution had the same effect in another background was the swap into AncB4 of mutations that evolved in AncB5– the most similar genotypes of all pairs of starting points in the analysis. Contingency therefore arose because historical substitutions that occurred during the intervals between ancestral proteins made specificity-changing mutations either deleterious or functionally inconsequential when introduced at times before or after the interval during which those mutations occurred.

**Figure 6.**
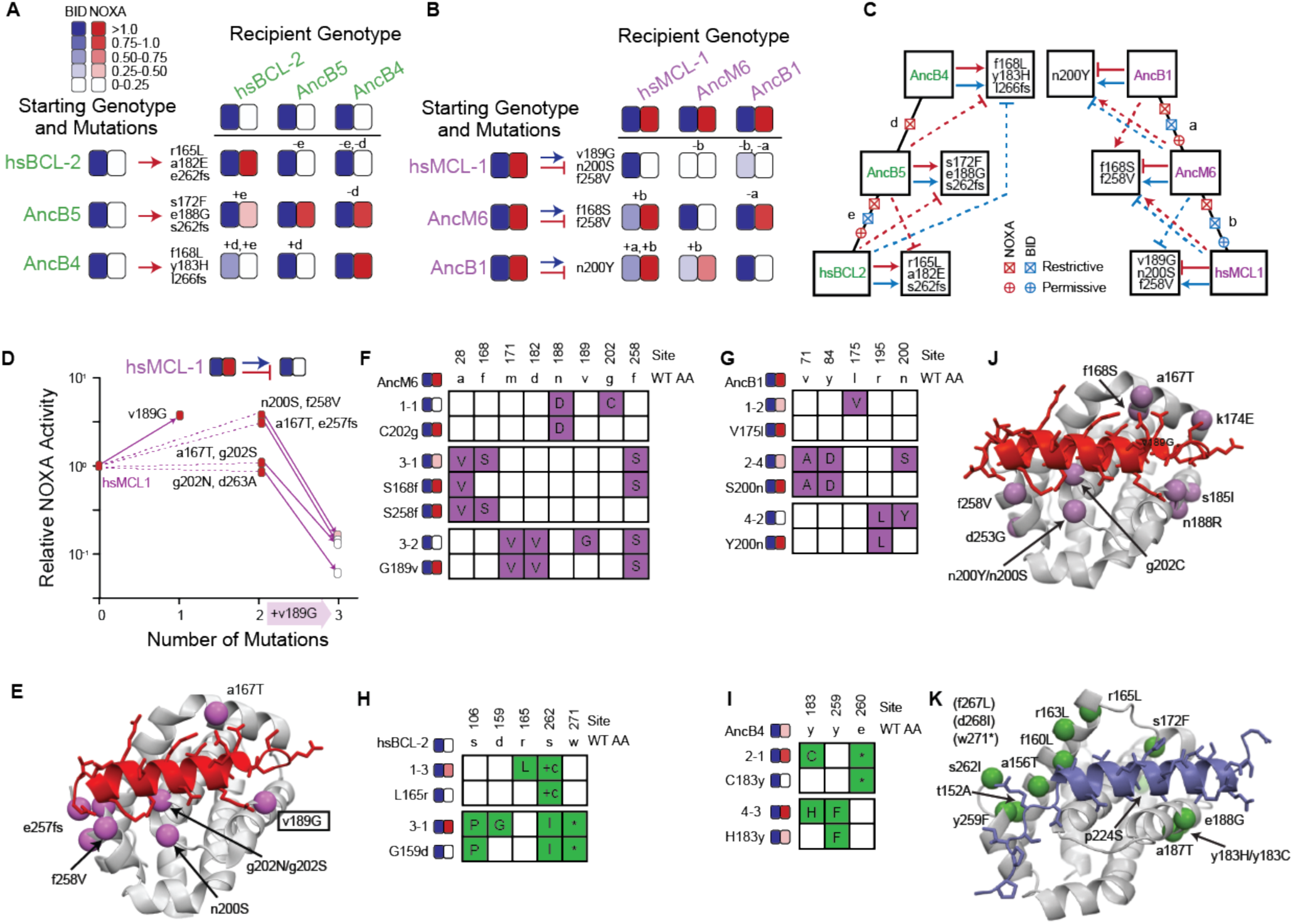
Sources of chance and contingency. (A) Epistatic incompatibility of PACE mutations in other historical proteins. Effects on activity are shown when amino acids states acquired in PACE under selection to acquire NOXA binding (red arrows) are introduced into ancestral and extant proteins. The listed mutations that occurred during PACE launched from each starting point (rows) were introduced as a group into the protein listed for each column. Observed BID (blue) and NOXA (red) activity in the luciferase assay for each mutant protein are shown as heatmaps (normalized mean of three biological replicates). Letters above each cell indicate the phylogenetic branch in Figure 3D that connects the PACE starting genotype to the recipient genotype for that experiments. Plus and minus signs indicate whether mutations were introduced into a descendant or more ancestral sequence, respectively. (B) Effects on activity when amino acids acquired in PACE under selection to lose NOXA binding and acquire BID binding are introduced into different ancestral and extant proteins, represented as in panel A. (C) Epistatic interactions between historical substitutions and PACE mutations. Restrictive historical substitutions (X) cause mutations that alter PPI specificity in an ancestor to abolish either BID (blue) or NOXA (red) activity when introduced into later historical proteins. Permissive substitutions (+) cause PACE mutations that alter PPI specificity in a descendent to abolish either BID or NOXA activity in an ancestor. Arrow, gain or maintenance of binding. Blunt bar, loss of binding. Mutations that confer selected functions in PACE are shown in the boxes at the end of solid arrows or bars. Solid lines, functional changes under PACE selection. Dashed lines, functional effects different from those selected for when PACE derived mutations are placed on a different genetic background. (D) Dissecting the effects of sets of mutations (white boxes) that caused hsMCL-1 to lose NOXA binding during four PACE trajectories. Filled boxes show the effect of introducing a subset of mutations into hsMCL-1 (normalized mean relative from three biological replicates). Solid lines show the effect of introducing v189G, which was found in all four sets. Dotted lines, effects of the other mutations in each set. (E) Structural location of mutations in panel D. Alpha-carbon atom of mutated residues are shown as purple spheres on the structure of MCL-1 (light gray) bound to NOXA (red, PDB 2nla). (F) Phenotypic effects of reverting frequent PACE-derived mutations when selecting AncM6 to lose NOXA activity. Individual variants were isolated from PACE experiments and sequenced. Sites and wild-type amino acid (WT AA) state are indicated at top. For each variant, non-WT states are shown in purple. Heatmaps on the left show binding to BID and NOXA in the luciferase assay for each variant and with a key mutation reverted back to the wild-type state corresponding mutant without the key mutation. Each shaded box represents the normalized mean of three biological replicates. (G) Phenotypic effects of reverting frequent PACE-derived mutations when selecting AncB1 to lose NOXA activity. (H) Phenotypic effects of reverting frequent PACE-derived mutations when selecting hsBCL-2 to gain NOXA activity. For each variant, non-WT states are shown in green. (I) Phenotypic effects of reverting frequent PACE-derived mutations when selecting AncB4 to lose NOXA activity. (J) Location of repeated mutations when hsMCL-1, AncM6, and AncB1 were selected to lose NOXA binding (purple spheres), represented on the structure of MCL-1 (gray) bound to NOXA (red, PDB 2nla). (K) Location of repeated mutations when hsBCL-2, AncB5, and AncB4 were selected to gain NOXA binding (green spheres), on the structure of hsBCL-xL (gray) bound to BID (blue, PDB 4qve).

To characterize the timing and effect of these epistatic substitutions during historical evolution, we mapped the observed incompatibilities onto the phylogeny (Figure 6C). We inferred that restrictive substitutions evolved on a branch if mutations that arose during directed evolution of an ancestral protein compromised coregulator binding when swapped into descendants of that branch. Conversely, we inferred that permissive substitutions evolved on a branch if mutations that arose during directed evolution compromised coregulator binding when swapped into more ancient ancestral proteins.

We found that both permissive and restrictive epistatic substitutions occurred on almost every branch of the phylogeny and affected both BID and NOXA binding. The only exception was the branch from AncB4 to AncB5, on which only restrictive substitutions affecting NOXA binding occurred. This is the branch immediately after NOXA function changed during history, the shortest of the branches examined, and the one with the smallest effect of contingency on genetic variance (Figure 5B). Even across this branch, the PACE mutations that restore the ancestral PPI specificity in AncB4 can no longer do so in AncB5. These results indicate that the paths through sequence space that could allow historical PPI specificities to be acquired repeatedly changed during the BCL-2 family’s history, even during intervals when the proteins’ PPI binding profiles did not change.

### Chance is caused by degeneracy in sequence-function relationships

For chance to strongly influence the outcomes of adaptive evolution, multiple paths to a selected phenotype must be accessible with similar probabilities of being taken. This situation could arise if several different mutations (or sets of mutations) can confer a new function, or if there are mutations that have no effect on function that can accompany function-changing mutations by chance. To distinguish between these possibilities, we measured the functional effect of different sets of mutations that arose in replicates when hsMCL-1 was evolved to lose NOXA binding (Figures 6D and S6A). One mutation (v189G) was found at high frequency in all four replicates, but it was always accompanied by other mutations, which varied among trajectories. We found that the apparently deterministic mutation v189G was a major contributor to the loss of NOXA binding, but it had this effect only in the presence of the other mutations, which did not decrease NOXA binding on their own. Mutation v189G therefore required permissive mutations to occur during directed evolution, and there were multiple sets of mutations with the potential to exert that effect; precisely which permissive mutations occurred in any replicate was a matter of chance. Despite this effect, all permissive mutations were located near the NOXA binding cleft, suggesting a shared mechanistic basis (Figure 6E).

Other starting genotypes showed a similar pattern of multiple mutations capable of conferring the selected function (Figures 6F-I, S6B, and S6C). In addition, when mapped onto the protein structure, all sites that were mutated in more than one replicate either directly contacted the bound peptide or were on secondary structural elements that did so (Figures 6J and 6K), suggesting a limited number of structural mechanisms by which PPIs can be altered. Taken together, these results indicate that chance arose because from each starting genotype, there were multiple mutational paths to the selected specificity; partial determinism arose because the number of accessible routes was limited by the structure-function relationships required for peptide binding in this family of proteins.

To more deeply understand the genetic basis for the limited degree of predictability that we observed, we performed PACE experiments in which we evolved hsBCL-2 to retain its BID binding, without selection for or against NOXA binding; we then screened for variants that fortuitously gained NOXA binding using an activity-dependent plaque assay (Figures S7A and S7B). This strategy allowed us to distinguish the number of mutations that can confer NOXA binding from the potentially more limited set of mutations with the highest selection coefficient(s). All four replicate populations produced clones that neutrally gained NOXA binding at a frequency of between ∼0.1% to 1% – lower than when NOXA binding was directly selected for, but five orders of magnitude higher than when NOXA binding was selected against (Figure 7A). From each replicate, we sequenced three NOXA-binding clones and found that all but one of them contained mutation r165L (Figure 7B), a mutation that also occurred at high frequency when the same protein was selected to gain NOXA binding. We introduced r165L into hsBCL-2 and found that it conferred significant NOXA binding with little effect on BID binding (Figure S7C). Several other mutations also appeared repeatedly in clones that fortuitously acquired NOXA binding, and these mutations were also acquired under selection for NOXA binding. A similar pattern of common mutations was observed in AncB4 and AncB5 clones that fortuitously or selectively evolved NOXA binding (Figures S7F-J). These observations indicate that the partial determinism we observed arises because from any specific starting genotype, there are only a few genotypes that can increase NOXA binding while retaining BID binding – not because there are many such genotypes, but under strong selection a few are strongly favored over others.

**Figure 7.**
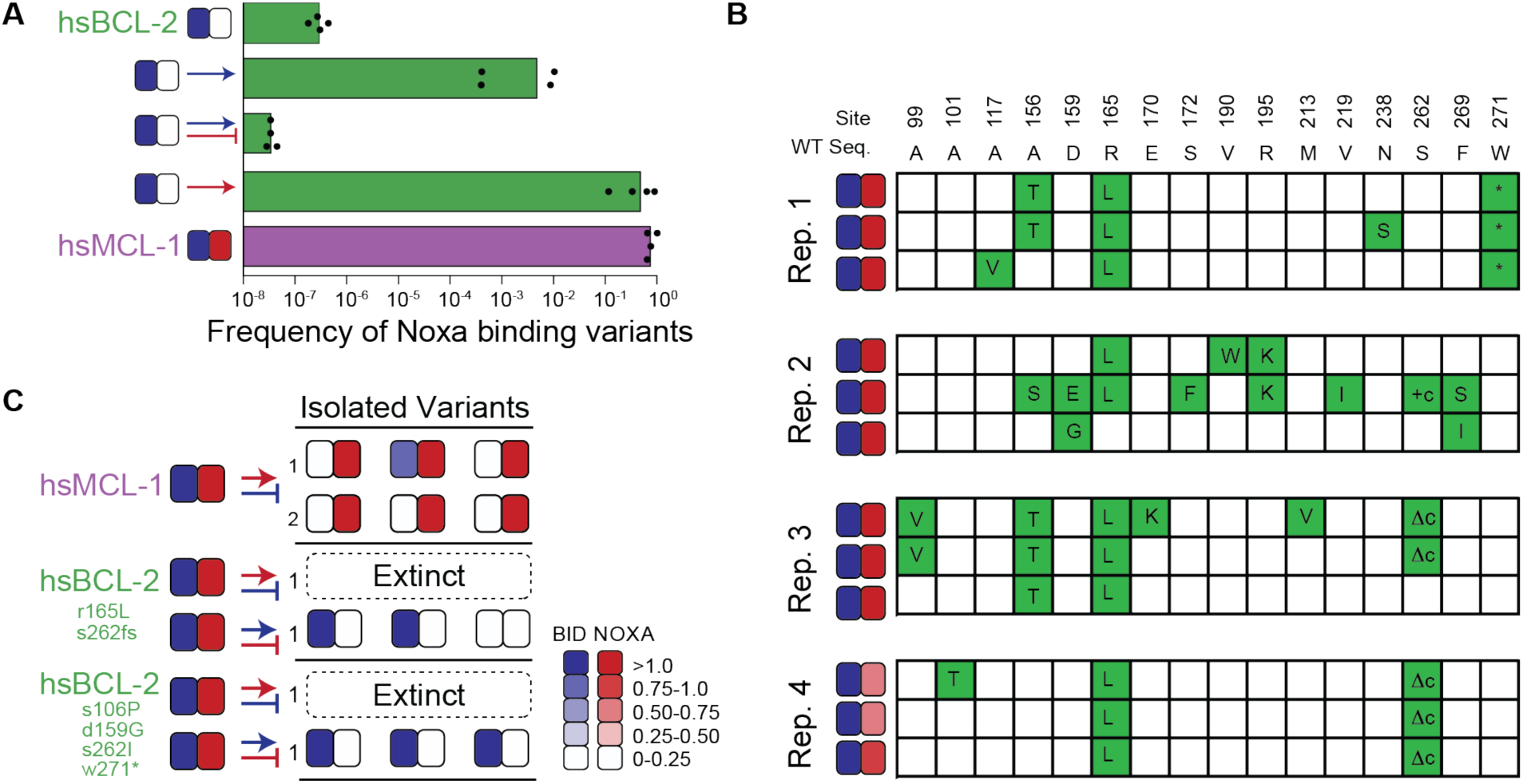
Evolutionary consequences of chance and contingency. (A) Evolution NOXA-binding phage from various selection regimes. Frequency was calculated as the ratio of plaque forming units (PFU) per milliliter on *E. coli* cells that require NOXA binding to the PFU on cells that require BID binding to form plaques. Wildtype hsBCL-2 (green) and hsMCL-1 (purple) are shown as controls. Arrow, positive selection for function. Bar, counter selection against function. Blue, BID. Red, NOXA. Columns show the mean of four trajectories under each condition (points). (B) Phenotypes and genotypes of hsBCL-2 variants that evolved NOXA binding under selection only to maintain BID binding. Sites and WT amino state are indicated at top. For each variant, non-WT states acquired are shown in green. Heatmaps show binding to BID and NOXA in the luciferase assay for each variant (normalized mean of three biological replicates). (C) Phenotypic outcomes when selecting for non-historical functions in PACE. Heatmaps show binding to BID and NOXA in the luciferase assay for each starting genotype (on the left) and for three individual variants picked from the trajectory’s outcome. Each shaded box shows the normalized mean of three biological replicates.

### The accessibility of new functions is affected by contingency

Although we found a strong influence of chance and contingency on the sequence outcomes of evolution across ancestral starting points, all replicates from all starting points acquired the selected BCL-2- or MCL-1-like specificity, indicating strong necessity at the level of protein function. This was true whether evolution began from more “promiscuous” starting points that bound both BID and NOXA or from more specific proteins that bound only BID. To further probe the evolutionary accessibility of new functions, we used PACE to select for a PPI specificity that never arose during historical evolution – binding of NOXA but not BID. We found that trajectories launched from hsMCL-1 (which binds both coregulators) readily evolved the selected phenotype, but two PACE-evolved variants of hsBCL-2, which had acquired the same PPI profile, went extinct under the same selection conditions (Figures 7C and S7K-M). The inability of the derived hsBCL-2 genotypes to acquire NOXA specificity was not attributable to a general lack of evolvability, because these same genotypes successfully evolved in a separate PACE experiment to lose their NOXA binding but retain BID binding (Figure S7N). These results establish that contingency can influence the accessibility of new functions during history and that apparently promiscuous proteins are not necessarily more able to evolve new PPI profiles than proteins that are more initially specific.

## DISCUSSION

The two major paradigms of 20^th^ Century evolutionary biology – the adaptationist program (Mayr 1983) and the neutral theory of molecular evolution (Kimura 1986) – focus on either necessity or chance, respectively, as the primary mode of causation that produces natural variation. Neither of these schools of thought admits much influence from contingency or history. From an adaptationist perspective, variation is generated by natural selection, which generates optimal forms under different environmental conditions. For example, differences in protein sequence, stability, or multimerization are interpreted as adaptive changes that improve a molecule’s ability to perform its function in the species’ particular environment (Somero 1995; Závodszky et al. 1998; Goodsell and Olson 2000; Nguyen et al. 2016). For neutralists, variation reflects the influence of chance in choosing among biologically equivalent possibilities, and conservation reflects purifying selection, both of which are largely unchanging across sequences in an alignment. For example, conserved portions of molecular sequences are interpreted as essential to structure and function, whereas differences in sequence alignments reflect a lack of constraint (Perutz et al. 1965; Kimura and Ota 1974; Echave et al. 2016). In neither worldview does the particular state of a biological system strongly reflect its past or shape its evolutionary future. Recent work has shown that contingency might be important in shaping the sequence outcomes of protein evolution (Ortlund et al. 2007; Blount et al. 2008, 2012; Bridgham et al. 2009; Bloom et al. 2010; Breen et al. 2012; Pollock et al. 2012; Shah et al. 2015; Sailer et al. 2017; Starr et al. 2018), echoing themes raised in paleontology (Gould 1990; Jablonski 2017) and developmental biology (Gompel et al. 2005; Shubin et al. 2009). Despite these recent findings, the dominance of the adaptationist and neutralist worldviews -- and the continuing rhetorical battle between them (Kern and Hahn 2018; Jensen et al. 2019)-- has obscured the possibility that contingency might join selection, drift, and mutation as a primary factor shaping the outcomes of evolution.

We found that contingency plays a profound role on phylogenetic timescales. The mutations that rose to high frequency during experimental evolution were almost completely different between individual evolutionary trajectories initiated from historical starting points separated by long phylogenetic distances. We observed a strong role for chance (because trajectories launched from the same starting point evolved extensive differences from each other) and an even greater effect of contingency (because pools of trajectories launched from different starting genotypes evolved even greater differences). When combined, chance and contingency erased virtually all traces of necessity between individual trajectories initiated from distantly related starting points. With the exception of a single truncation mutation that does not affect the selected-for function, the only predictable sequence states were those that were unchanged from the starting point in all trajectories, presumably because they are unconditionally necessary for the historically relevant PPIs tested and were therefore conserved by purifying selection.

In quantitative terms, chance and contingency multiplied each other’s effects, magnifying the probability that evolutionary trajectories launched from different starting points across history will lead to different sequence outcomes. This relationship is a feature of the way genetic variance is analyzed, but it also reflects the qualitative interactions between chance and contingency across time. At every point in history, there were numerous sets of mutations with the potential to confer a selected-for PPI specificity, and chance determined which was taken. These chance steps then determined the steps that can be taken in the next and future intervals. Without contingency, chance events would be inconsequential, because every path that was ever open would remain forever so, irrespective of the random events that happen to take place. Conversely, without chance, contingency in history would be inconsequential, because all phylogenetic lineages launched from a common ancestor would always lead to the same intermediate steps and thus the same ultimate outcomes. Thus, the outcomes of evolution are unpredictable because intermediate steps shape future possibilities, and those intermediate steps cannot be predicted because multiple routes are available to evolution at any point in history.

Our experimental design comes close to, but does not quite achieve, the ideal design of multireplicate evolution from ancestral starting points under historical conditions, because the particular selection pressures, environments, and population parameters that pertained in the deep past are unknown. It is likely, however, that chance and contingency were at least as significant during history as in our experiments. The relative roles of chance, contingency, and necessity are determined by population genetic parameters and by internal factors that arise from the protein’s physical architecture. The population genetic parameters in our experimental conditions are likely to strongly favor determinism, because they involve very large population sizes, strong selection pressures, and high mutation rates, all directed at a single gene. If historical evolution in the BCL-2 family involved smaller populations, weaker selection, lower mutation rates, and a larger genetic “target size” for adaptation, as seems likely, then chance would have played an even larger role during history than in our experiments. Second, chance and contingency arise from the relationship between a protein’s sequence, structure, and function, such as the number of mutations that can produce a particular function and the nature of epistasis among them. Tertiary structure is highly conserved across BCL-2 family members, and they bind coregulators using the same surface. It is therefore unlikely that the internal determinants of chance, contingency, and necessity varied strongly between history and our experimental regime.

Although we studied the BCL-2 family as a model, we expect that our qualitative results will apply to many other proteins. Epistasis is a common feature of protein structure and function, so the accumulating effect of contingency across phylogenetic time in the BCL-2 family will probably be a general feature of protein evolution, although its rate and extent are likely to vary among protein families and timescales (Chandler et al. 2013; Harms and Thornton 2014; Shah et al. 2015; Zhu et al. 2018). The influence of chance depends upon the existence of multiple mutational sets that can confer a new function; this kind of degeneracy is likely to pertain in many cases: greater determinism is expected for functions with very narrow sequence-structure-function-constraints, such as catalysis (Menéndez-Arias 2010; Salverda et al. 2011; Meyer et al. 2012; Storz 2016; Hawkins et al. 2019; Karageorgi et al. 2019), than those for which sequence requirements are less strict, such as substrate binding (Yokoyama et al. 2008; Blount et al. 2012; Starr et al. 2017; Zheng et al. 2019). Consistent with this prediction, when experimental evolution regimes have imposed diffuse selection pressures on whole organisms, making loci across the entire genome potential sources of adaptive mutations, virtually no repeatability has been observed among replicates (Kryazhimskiy et al. 2014; Wünsche et al. 2017).

Our results have implications for efforts to engineer proteins with desired properties. We found no evidence that ancestral proteins were more or less “evolvable” than extant proteins: the selected-for phenotypes readily evolved from both extant and ancestral proteins with the same starting binding capabilities. Moreover, chance’s effect was virtually constant across ∼ 1 billion years of evolution, indicating that the number of accessible mutations in the deep past that could confer a selected-for function was apparently no greater than it is now. Nevertheless, the strong effect of contingency suggests that efforts to produce proteins with new functions by design or directed evolution will be most effective if they use multiple different protein sequences as starting points, ideally separated by long intervals of sequence evolution. Ancestral proteins can be useful for this purpose, simply because the particular routes to new functions they provide are different from those provided by any extant protein, even if those routes are not fundamentally different in number or kind.

The method that we developed for rapid evolution of PPI specificity has several distinct advantages that can be extended to other protein families. First, by using PACE, many replicates can be evolved in parallel across scores or hundreds of generations in just days, with minimal need for intervention by the experimentalist (Esvelt et al. 2011). Second, our split RNA-polymerase design for acquiring new PPIs has fewer components than previous methods for this purpose, such as two-hybrid designs; this makes it considerably easier to tune and optimize and therefore to extend to other protein systems. Third, unlike approaches that attempt to evolve specific PPIs by alternating selection and counterselection through time, our platform simultaneously imposes selection and counterselection within the same cell, thus selecting for specificity directly. By combining these elements in a single system, our platform should allow rapid multireplicate evolution of new PPI specificities in a variety of protein families.

Finally, our work has implications for the processes of protein evolution and the significance of natural sequence variation. Our findings suggest that sequence-structure-function associations apparent in sequence alignments are, to a significant degree, the result of contingent constraints that were transiently imposed or removed by chance events during history (Gong et al. 2013; Harms and Thornton 2014; Starr et al. 2017, 2018). Evolutionary explanations of sequence diversity and conservation must therefore explicitly consider the historical trajectories by which sequences evolved, in contrast to the largely history-free approaches of the dominant schools of thought in molecular evolution. Present-day proteins are physical anecdotes of their particular unpredictable histories: their sequences reflect the interaction of accumulated chance events during descent from common ancestors with necessity imposed by physics, chemistry, and natural selection. Apparent “design principles” in the pattern of variability and conservation in extant proteins reflect not how things must be to perform their functions, or even how they can best do so; rather, today’s proteins reflect the legacy of the opportunities and limitations that they just happen to have inherited.

## MATERIALS AND METHODS

### Phylogenetics

Amino acid sequences of the human BCL-2, BCLW, BCL-xL, MCL-1, NRH, BFL1, BAK, BAX, and BOK paralogs were used as starting points for identifying BCL-2 family members in other species. For each paralog, tblastn and protein BLAST on NCBI BLAST were used to identify orthologous sequences between January and March of 2018 (Altschul et al. 1997). Sequences for each paralog were aligned using MAFFT (G-INS-I) with the --allowshift option and -- unalignlevel set at 0.1. For each paralog, phylogenetic structure was determined using fasttree 2.1.11 within Geneious 10.1.3. Missing clades based on known species relationships were then identified and specific tblastn searches were used within Afrotheria (taxid:311790), Marsupials (taxid:9263), Monotremes (taxid:9255), Squamata (taxid:8509), Archosauria (taxid:8492), Testudinata (taxid:8459), Amphibia (taxid:8292), Chondrichthyes (taxid:7777), Actinopterygii (taxid:7898), Dipnomorpha (taxid:7878), Actinistia (taxid:118072), Agnatha (taxid:1476529), Cephalochordata (taxid:7735), and Tunicata (taxid:7712) as needed. Additional sequences were added by downloading genome and transcriptome data for tuatara (Miller et al. 2012), sharks and rays (Wyffels et al. 2014), gar (Zerbino et al. 2018), ray-finned fish (Hughes et al. 2018), lamprey (Smith et al. 2018), hagfish (Takechi et al. 2011), *Ciona savignyi* (Zerbino et al. 2018), tunicates (Delsuc et al. 2018), echinoderms (Reich et al. 2015), porifera (Riesgo et al. 2014), and ctenophores (Moroz et al. 2014). In each case, local BLAST databases were created in Geneious and searched using tblastn. Finally, we used BCL-2DB to add missing groups as needed (De Laval et al. 2014).

After collection of sequences, each paralog was realigned using MAFFT (G-INS-I) with the --allowshift option and --unalignlevel set at 0.1. Based on known species relationships, lineage specific insertions were removed and gaps manually edited. Only a single sequence was kept among pairs of sequences differing by a single amino acid and sequences with more than 25% of missing sites were removed. For difficult to align sequences, sequences were modeled on the structures of human BCL-2 family members using SWISS-Model to identify likely locations of gaps (Waterhouse et al. 2018). Finally, paralogs were profile aligned to each other and paralog specific insertions were identified.

In total, 151 amino acid sites from 745 taxa were used to infer the phylogenetic relationships among BCL-2 paralogs. PROT Test 3.4.2 was used to identify the best fit model among JTT, LG, and WAG, with combinations of observed amino acid frequencies (+F), gamma distributed rate categories (+G), and an invariant category (+I). From this, JTT + G + F had the highest likelihood and lowest AIC. RAXML-ng 0.6.0 was then used to identify the maximum likelihood tree using JTT+G12+F0 (12 gamma rate categories with maximum likelihood estimated amino acid frequencies) (Kozlov and Stamatakis 2019). Finally, we enforced monophyly within each paralog for the following groups: lobe-finned fish (n=9), ray-finned fish (n=9), jawless fish (n=5), cartilaginous fish (n=8), tunicates (n=4), branchiostoma (n=4), chordates (n=5), ambulacraria (n=5, hemichordata + echinodermata), deuterostomia (n=5), protostomia (n=5), cnidaria (n=5), and porifera (n=4) (values in parenthesis are number of identified paralogs in each group) and used RAXML-ng with JTT + G12 + F0 to identify the best tree given these constraints (Supplementary Data Phylogenetic.Data.zip).

Overall, we recovered three clades; a pro-apoptotic clade, a clade containing the BCL-2, BCLW, and BCLX paralogs, and a clade containing the MCL-1, BFL1, and NRH paralogs. We used the pro-apoptotic clade as the outgroup to the two anti-apoptotic clades. Within the BCL-2 clade, the majority of vertebrates contained all three copies. However, the exact relationship among the paralogs was unclear; only two copies were identified within jawless fish and their phylogenetic placement had weak support. Non-vertebrate clades tended to have good support and only a single copy. However, support for these groups following established species relationships was often limited. The MCL-1 clade contained the fastest evolving paralogs of the BCL-2 family. As with the BCL-2-like clade, only two copies were found within the jawless fish and the exact sister relationships among paralogs was unclear. Non-vertebrates contained only a single copy, but as with the BCL-2-like clade, support for relationships following established species relationships was often weak.

The BCL-2-like and MCL-1-like paralogs formed a clade with the BHP1 and BHP2 sequences from porifera. The sister relationships among these four clades were unresolved. In addition, we recovered a sister relationship between the BAK and BAX paralogs. While both paralogs contained copies from porifera, these clades evolved quickly and had relatively low support, and they may be artifactual. We identified only a single clade of ctenophores. Finally, the placement of BOK was unresolved; BOK may be sister to the BAK/BAX clade or an outgroup to all clades and the most ancient copy of the BCL-2 family.

### Ancestral reconstruction

Posterior probabilities of each amino acid at each site were inferred using Lazarus (Finnigan et al. 2012) to run codeml within PAML. We used the same model and alignment as used to infer the phylogeny. We used the branch lengths and topology of the constrained maximum likelihood phylogeny found by raxml-ng.

We first reconstructed the last common ancestors (LCA) of all BCL-2 and MCL-1 like sequences, AncMB1-M, using the maximum likelihood state for each alignable site. We then reconstructed a series of ancestors from AncMB1 to modern human MCL-1. These included AncM1 (last common ancestor of MCL-1 related sequences), AncM2 (LCA of MCL-1 related Deuterostomes and Protostomes), AncM3 (LCA of MCL-1 related Deuterostomes), AncM4 (LCA of MCL-1 related Urochordates and Chordates), AncM5 (LCA of MCL-1, BFL1, and NRH like copies in vertebrates), AncM6 (LCA of MCL-1 and BFL1 like copies), AncMCL-1 (LCA of MCL-1 like copies), AncMCL-1-G (LCA of MCL-1 like Gnathostomes), AncMCL-1-O (LCA of MCL-1 like Osteichthyes), and AncMCL-1-T (LCA of MCL-1 like Tetrapods), AncMCL-1-A (LCA of MCL-1 like Amniotes), and AncMCL-1-M (LCA of MCL-1 like Mammals). In each case, the sequence of each ancestor used the maximum likelihood state at each site, with gaps inserted based on parsimony. We used the modern sequences of human MCL-1 to fill in portions of the sequence that showed poor alignment and could not be reconstructed, including both the N and C-terms, as well as the loop between the first and second alpha-helices. Average posterior probabilities for ancestors in the MCL-1 clade ranged from 0.73 (AncM6) to 0.98 (AncMCL-1-M) with an average of 0.83 (sd 0.08) (Table S2).

For the BCL-2 like clade, we also reconstructed AncMB1, this time using human BCL-2 sequence to fill in the N and C-terms and the loop between the first and second alpha helices (AncMB1-B). We then reconstructed sequences from AncMB1 to modern human BCL-2. These included AncB1 (last common ancestor of BCL-2 related sequences), AncB2 (LCA of BCL-2 related Bilaterian and Cnidaria), AncB3 (LCA of BCL-2 related Deuterostomes and Protostomes), AncB4 (LCA of BCL-2 Deuterostomes), AncB5 (LCA of BCL-2, BCLW, and BCLX like copies in vertebrates), AncBCL-2 (LCA of BCL-2 like copies), AncBCL-2-G (LCA of BCL-2 like Gnathostomes), AncBCL-2-O (LCA of BCL-2 like Osteichthyes), and AncBCL-2-T (LCA of BCL-2 like Tetrapods), using human BCL-2 sequences for the N and C-terms and the loop between the first and second alpha helices. Average posterior probabilities for ancestors in the BCL-2 clade ranged from 0.87 (AncB1) to 0.95 (AncBCL-2-T) with an average of 0.9 (sd 0.04).

### Test of robustness of ancestral inference

To determine the robustness of our conclusions on the phenotype of ancestral sequences, we synthesized and cloned alternative reconstructions for key ancestors. In each case, sequences contained the most likely alternative state with posterior probability > 0.2 for all such sites where such a state existed. Alternative reconstructions contained an average of 24 alternative states and represent a conservative test of function (min: 4, max: 44, Table S2). In our luciferase assay, all but two alternative reconstructions retained similar BID and NOXA binding as the maximum likelihood ancestral sequences. The first alternative reconstruction that differed from the ML reconstruction was AltAncB3, which bound both BID and NOXA, while the ML for AncB3 bound BID, but NOXA only weakly. As a result, the exact branch upon which NOXA binding was lost historically is not resolved by this data.

The second alternative reconstruction that differed from the ML reconstruction was AltAncMB1-B, which had weaker NOXA binding than the ML reconstruction. To further test the robustness of AncMB1-B to alternative reconstructions, we synthesized and tested additional reconstructions that included only alternative amino acids with posterior probabilities greater than 0.4 (n=3), 0.35 (n=7), 0.3 (n=13), and 0.25 (n=18), and compared these to AncMB1-B and the 0.2 AltAncMB1-B (n=21) (values in parentheses are number of states that differ from the ML state). We found that the 0.4, 0.35, and 0.3 alternative reconstructions bound both BID and NOXA, while the 0.25 and 0.2 alternative reconstructions had diminished NOXA binding.

Finally, we synthesized and tested modern sequences from key groups to determine the robustness of our inference on the timing of NOXA binding loss. These included BCL-2 related sequences from groups that diverged prior to the predicted loss of NOXA binding (*Trichoplax adhaerens* and *Hydra magnapapillata*), sequences from groups that diverged around the time of predicted NOXA binding loss (*Octopus bimaculoides* and *Stegodyphus mimosarum*), or sequences from groups predicted to have diverged after NOXA binding lost (*Saccoglossus kowalevskii* and *Branchiostoma belcheri*). While the *T. adhaerens* and *B. belcheri* sequences were non-functional in our luciferase assays, binding neither BID nor NOXA, *H. magnapapillata* bound both BID and NOXA and the remaining sequences bound only BID, suggesting a loss of NOXA binding prior to the divergence of protostomes and deuterostomes in the BCL-2 related clade, consistent with the conclusion drawn using reconstructed proteins.

### *Escherichia coli* strains

*E. coli* 10-beta cells were used for cloning and were cultured in 2xYT media. *E. coli* BL21 (BE3) cells were used for protein expression and were cultured in Luria-Bertain (LB) broth. *E. coli* S1030 cells cultured in LB broth were used for activity-dependent plaque assays, phage growth assays, and luciferase assays. S1030 cells cultured in Davis Rich media were used for PACE experiments (Carlson et al. 2014). *E. coli* 1059 cells were used for cloning phage and assessing phage titers and were cultured in 2xYT media

### Cloning and general methods

Plasmids were constructed by using Q5 DNA Polymerase (NEB) to amplify fragments that were then ligated via Gibson Assembly. Primers were obtained from IDT, and all plasmids were sequenced at the University of Chicago Comprehensive Cancer Center DNA Sequencing and Genotyping Facility. Vectors and gene sequences used in this study are listed in Table S5, with links to fully annotated vector maps on Benchling. Key vectors are deposited at Addgene, and all vectors are available upon request. The following working concentrations of antibiotics were used: 50 µg/mL carbenicillin, 50 µg/mL spectinomycin, 40 µg/mL kanamycin, and 33 µg/mL chloramphenicol. Protein structures and alignments were generated using the program PyMOL (Schrödinger 2018).

### Luciferase assay

Cloned expression vectors contained the following: 1) a previously-evolved, isopropyl β-D-1-thiogalactopyranoside (IPTG)-inducible N-terminal half of T7 RNAP (Zinkus-Boltz et al. 2019) fused to a BCL-2 family protein; 2) the C-terminal half of T7 RNAP fused to a peptide from a BH3-only protein; and 3) T7 promoter-driven luciferase reporter. Chemically-competent S1030 *E. coli* cells (Carlson et al. 2014) were prepared by culturing to an OD_600_ of 0.3, washing twice with a calcium chloride/HEPES solution (60 mM CaCl_2_, 10 mM HEPES pH 7.0, 15% glycerol), and then resuspending in the same solution. Vectors were transformed into chemically-competent S1030 cells via heat shock at 42 ºC for 45 sec, followed by 1 h recovery in 3x volume of 2xYT media, and then plated on agar with the appropriate antibiotics (carbenicillin, spectinomycin, and chloramphenicol) to incubate overnight at 37 ºC. Individual colonies (3-4 biological replicates per condition) were picked and cultured in 1 mL of LB media containing the appropriate antibiotics overnight at 37 ºC in a shaker. The next morning, 50 µL of each culture was diluted into 450 µL of fresh LB media containing the appropriate antibiotics, as well as 1 µM of IPTG. The cells were incubated in a shaker at 37 ºC, and OD_600_ and luminescence measurements were recorded between 2.5 and 4.5 hours after the start of the incubation. Measurements were taken on a Synergy Neo2 Microplate Reader (BioTek) by transferring 150 µL of the daytime cultures into Corning^®^ black, clear-bottom 96-well plates. Data were analyzed in Microsoft Excel and plotted in GraphPad Prism, as previously reported (Pu et al. 2017a).

### Protein expression

hsBCL-2, hsMCL-1, and evolved variants were constructed as N-terminal 6xHis-GST tagged proteins. The recombinant proteins were expressed in BL21 *E. coli* (NEB) and purified following standard Ni-NTA resin purification protocols (ThermoFisher Scientific) (Zhou et al. 2019). Briefly, BL21 *E. coli* containing an N-terminal 6xHis-GST tagged BCL-2 family protein were cultured in 5 mL LB with carbenicillin overnight. The following day, the culture was added to 0.5 L of LB with carbenicillin, incubated at 37 ⍛C until it reached an OD_600_ of 0.6, induced with IPTG (final concentration: 200 µM), and cultured overnight at 16 ⍛C. The cell pellet was harvested by centrifugation followed by resuspension in 30 mL of lysis buffer (50 mM Tris 1 M NaCl, 20% glycerol, 10 mM TCEP, pH 7.5) supplemented by protease inhibitors (200 nM Aprotinin, 10 µM Bestatin, 20 µM E-64, 100 µM Leupeptin, 1 mM AEBSF, 20 µM Pepstatin A). Cells were lysed via sonication and were then centrifuged at 12,000 g for 40 min at 4 ⍛C. Solubilized proteins, located in the supernatant, were incubated with His60 Ni Superflow Resin (Takara) for 1 hr at 4 ⍛C, and the protein was eluted using a gradient of imidazole in lysis buffer (50-250 mM). Fractions with the protein, as determined by SDS-PAGE, were concentrated in Ulta-50 Centrifugal Filter Units (Amicon, EMD Millipore). Proteins were purified via a desalting column with storage buffer (50mM Tris-HCl (pH 7.5), 300 mM NaCl, 10% glycerol, 1 mM DTT) and further concentrated. The concentration of the purified BCL-2 family proteins was determined by BCA assay (ThermoFisher Scientific), and they were flash-frozen in liquid nitrogen and stored at -80⍛C.

### Fluorescent polarization binding assays

Fluorescent polarization (FP) was used to measure the affinity of BCL-2 family proteins with peptide fragments of the BH3-only proteins in accordance with previously-described methods (Zhang et al. 2002). hsBCL-2, hsMCL-1, and evolved variants were purified as described above. The fluorescent NOXA and BID peptides (95+% purity) were synthesized by GenScript and were N-terminally labeled with 5-FAM-Ahx and C-terminally modified by amidation. These peptides were dissolved and stored in DMSO. Corning^®^ black, clear-bottom 384-well plates were used to measure FP, and three replicates were prepared for each data point. Each well contained the following 100 µL reaction: 20 nM BH3-only protein, 0.05 nM to 3 µM of BCL-2 family protein (1/3 serial dilutions), 20 mM Tris (pH 7.5), 100 mM NaCl, 1 mM EDTA, and 0.05% pluronic F-68. FP values (in milli-polarization units; mFP) of each sample were read by a Synergy Neo2 Microplate Reader (BioTek) with the FP 108 filter (485/530) at room temperature 5-15 minutes after mixing all the components. Data were analyzed in GraphPad Prism 8, using the following customized fitting equation, to calculate *K*_*d*_(Zhou et al. 2019):

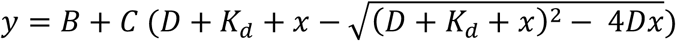

where *y* is normalized measured FP, *x* is the concentration of BCL-2 protein, *D* is the concentration of the BH3-only protein, *B* and *C* are parameters related to the FP value of free and bound BH3-only protein, and *K*_*d*_ is the dissociation constant.

### Phage-assisted continuous evolution

Phage-assisted continuous evolution (PACE) was used to evolve hsBCL-2, hsMCL-1, and ancestral proteins in accord with previously reported technical methods (Esvelt et al. 2011; Carlson et al. 2014; Pu et al. 2017b, 2019) using a new vector system. Briefly, combinations of accessory plasmids and the MP6 mutagenesis plasmid (Badran and Liu 2015) were transformed into S1030 *E. coli*., plated on agar containing the appropriate antibiotics (carbenicillin, kanamycin, and chloramphenicol) and 10 mM glucose, and incubated overnight at 37 ⍛C. Colonies were grown overnight in 5 mL of LB containing the appropriate antibiotics and 20 mM glucose. Davis Rich media was prepared in 5-10 L carboys and autoclaved, and the PACE flasks and corresponding pump tubing were autoclaved as well. The following day, PACE was set up in a 37 ⍛C environmental chamber (Forma 3960 environmental chamber, ThermoFisher Scientific). For each replicate, an overnight culture was added to ∼150 mL of Davis Rich carboy media in chemostats and grown for 2-3 h until reaching an OD_600_ of approximately 0.6. Lagoons containing 20 µL of phage from saturated phage stocks (10^8^-10^9^ phage) were then connected to the chemostat. Magnetic stir bars were used to agitate chemostats and lagoons. The chemostat cultures were flowed into the lagoons at a rate of approximately 20 mL/h. Waste output flow rates were adjusted to maintain a constant volume of 20 mL in the lagoons, 150 mL in the chemostat, and an OD_600_ close to 0.6 in the chemostat. A 10% w/v arabinose solution was pumped into the lagoons at a rate of 1 mL/h. If the experiment included a mixing step (two separate chemostats flowed together into one lagoon for a mixed selection pressure), a chemostat was prepared the next day (as described above) and connected to the lagoons. During this step, lagoon volumes were increased to 40 mL, and the arabinose inflow rate was increased to 2 mL/h. After disconnecting the first chemostat the next day, the lagoon volumes and arabinose inflow were both lowered to 20 mL and 1 mL/h, respectively. During the experiment, samples were collected from the lagoons every 24 h and centrifuged at 13,000 rpm for 3 min to collect the phage-containing supernatant, as well as the cell pellet for DNA extraction. PACE experiments are listed in Table S3.

During PACE, the media volume of each lagoon turned over once per hour for four days, or ∼100 times. For a phage population to survive this amount of dilution, a similar number of generations must have occurred between the starting phage and the phage in the lagoon at the end of the experiment (Esvelt et al. 2011). This is expected to be a conservative estimate; as a more fit phage rises in frequency in the population it will undergo a greater number of generations than less fit phage in the population. The mutagenesis plasmid MP6 induces a mutation rate of approximately 6×10^−6^ per bp per generation. The BCL2 family proteins used in the PACE experiments were ∼230 amino acids long, indicating that a mutation occurred on average every ∼250 phage replications. Phage population sizes ranged from 10^5^ per ml to 10^10^ per ml over the course of a PACE experiment, indicating a rate of 400 to 40,000,000 new mutations every generation. Conservative estimates thus suggest that a during each individual replicate, phage populations sampled at least 40,000 mutations, and upwards of 4×10^9^ mutations. While not all mutations were equally likely each generation because MP6 enriches for transitions (i.e. G⟶A, A⟶G, C⟶T, and T⟶C), the high number of mutations sampled suggests that the vast majority of possible single point mutations (approximately 230∗3∗4= 2760 potential mutations) were sampled over the course of each experiment, with higher population sizes generating all potential single point mutations each generation.

### Plaque assays

Plaque assays were performed on 1059 *E. coli* cells (Carlson et al. 2014; Hubbard et al. 2015), which supply gene III (gIII) to phage in an activity-independent manner, to measure phage titers. Additionally, activity-dependent plaque assays were done on S1030 *E. coli* containing the desired accessory plasmids to determine the number of phage encoding a BCL-2 family protein with a given peptide binding profile. All cells were grown to an OD_600_ of approximately 0.6 during the day. Four serial dilutions were done in Eppendorf tubes by serially pipetting 1 μL of phage into 50 µL of cells to yield the following dilutions: 1/50, 1/2500, 1/125000, and 1/6250000. 650 µL of top agar (0.7% agar with LB media) was added to each tube, which was then immediately spread onto a quad plate containing bottom agar (1.5% agar with LB media). Plates were incubated overnight at 37 ⍛C. Plaques were counted the following day, and plaque forming units (PFU) per mL was calculated using the following equation:

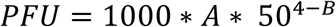

Where *A* is the number of plaques in a given quadrant, and *B* is the quadrant number where the phage were counted, in which 1 is the least dilute quadrant and 4 is the most dilute quadrant.

### Phage growth assays

Phage growth assays were performed by adding the following to a culture tube and shaking at 37 ⍛C for 6 h: 1 mL of LB with the appropriate antibiotics (carbenicillin and kanamycin), 10 µL of saturated S1030 *E. coli* containing the accessory plasmids of interest, and ∼1000 phage. Phage were then isolated by centrifugation at 13,000 rpm for 3 min, and PFU was determined by plaque assays using 1059 *E. coli* and the plaque assay protocol described above.

### High-throughput sequencing library construction

PACE samples were collected from each lagoon every 24 hours. The lagoon samples were centrifuged at 13,000 rpm for 3 min on a bench top centrifuge to separate supernatant and cell pellet. The phage-containing supernatants were stored at 4 ⍛C prior to the creation of sequencing libraries. To prepare Illumina sequencing libraries, each phage sample was cultured overnight with 1059 *E. coli* cells, followed by phage DNA purification (Qiagen plasmid purification reagents buffer P1 (catalog number 19051), P2 (catalog number 19052), N3 (catalog number 19064), PE (catalog number 19065), and spin column for DNA (EconoSpin™, catalog number 1920-250). The resulting DNA concentration was ∼50 ng/µL. Freshly generated DNA samples were then used as template for PCR amplification. For each library sample, we amplified three overlapping fragments of the BCL-2 family protein, which are 218∼241bp in length (Figure S4A). Each primer also included 6-9 “N”s to introduce length variation (Table S4). In total, 12 PCR reactions were used for each library. Phusion DNA polymerases and buffers (ThermoFisher Scientific, catalog number F518L) were used in the first PCR round to amplify all three fragments for all library sequencing. The 25 <u>μ</u>L reaction contained: 0.5 µL of 50 mM MgCl_2_, 0.75 µL of 10 mM dNTP, 0.75 µL Phusion DNA polymerase, 20 ng library DNA, 0.5 µL of 10µM primer (each). The PCR reactions were run on a C1000 Touch Thermal Cycler (Bio-Rad), with the following parameters: 98°C for 1 min, followed by 16 cycles of 98°C for 12 sec, 58°C for 15 sec and 72°C for 45 sec, and finally 72°C for 5 min. PCR reactions were purified using the ZYMO DNA clean and concentrator kit (catalog number D4013) and 96 well filter plate (EconoSpin, catalog number 2020-001). The DNA products were dissolved in 30 µL ddH_2_O. All 12 reactions for each library were combined and 1 µL was used as the template for a second PCR round. PCR reaction components and thermocycler parameters were the same as above, except that the annealing temperature was 56 °C, and only 15 rounds of amplification were conducted. The primer and sample combinations are listed in Table S4. PCR reactions were then purified following the same procedure as previous step. Equal volumes of all 72 library samples were combined and concentration was measured using a Qubit 4 Fluorometer. The total DNA sample was 2.68 ng/µL (equivalent to 10 nM, according to the average length of PCR fragments). DNA samples were diluted to 4 nM from step 4 following the Illumina MiSeq System Denature and Dilute Libraries Guide and then diluted to 12 pM for high-throughput sequencing. The final sample contained 100 µL of 20 pM PhiX spike-in plus 500 µL of the 12 pM library sample. Sequencing was performed on the Illumina MiSeq System using MiSeq Reagent Kit v3 (600-cycle) with paired-end reads according to the manufacturer’s instructions.

### Processing of Illumina data

Illumina sequencing yielded 22 million reads, 13 million of which could be matched to a specific sample (Table S4). One replicate for AncB5 was found to be contaminated and removed from further analysis. To process the remaining data, we first used Trim Galore with default settings to trim reads based on quality (https://www.bioinformatics.babraham.ac.uk/projects/trim_galore/). Then, we used BBMerge, a script in BBTools (https://jgi.doe.gov/data-and-tools/bbtools/), to merge paired-end reads. Next, we used Clumpify to remove repeated barcode sequences. We then used Seal to identify and bin reads by sample and fragment. Finally, we used BBDuk to remove any primer or adapter sequence present. Scripts and reference sequences are available on Github (https://github.com/JoeThorntonLab/BCL2.ChanceAndContingency).

### Illumina Sequencing Analysis

Reads were binned by experiment and then aligned to the appropriate WT sequence using Geneious (low Sensitivity, 5 iterations, gaps allowed). Sequences were then processed in R to remove sequences containing “N”s or that were not full length. Insertions found in less than 1% of the population and sites that extended outside of the coding region were removed from all sequences. Remaining gaps were standardized among replicates and within an experiment. Finally, allele frequencies were calculated for each site and amino acid, as well as remaining insertions and deletions.

### Quantifying the effects of chance and contingency on the outcomes of evolution

#### Estimating the effects of chance

Allele frequency differences among replicates started from the same genotype can only be caused by chance events. Thus, to determine the effects of chance (*C*_1_) on the outcomes of evolution, we compared allele frequencies from replicate PACE experiments started from the same genotype. We compared allele frequencies of individual replicates to the average allele frequencies among replicates started from the same genotype by estimating the probability that two randomly chosen alleles would be different, i.e. the genetic variance, for each replicate individually (*V*_*r*_) and the pooled sample of all replicates from a given starting genotype (*V*_*g*_):

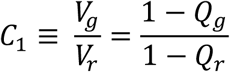

where *Q*_*r*_ is the probability that two randomly chosen alleles in the same replicate are identical in state and *Q*_*g*_ is the probability that two randomly chosen alleles from the pooled replicates started from the same genotype are identical in state. *C*_1_ is related to Wright’s F_is_ statistic as:

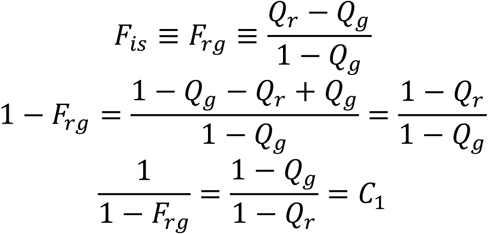

We used count data from Illumina sequencing to estimate allele frequencies and followed the approach of Hivert *et al*. 2018, which developed a methods of moments estimator,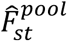, that is appropriate for pooled data and accounts for both the sampling of individuals within a population and the sampling of reads during sequencing. We treated each amino acid site independently and defined the following:

*R*_1:*rgs*_ ≡ # of reads in replicate *r* of starting genotype *g* at site *s*

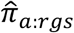 observed allele frequency of allele *a* in replicate *r* of starting genotype *g* at site *s*

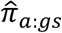 ≡ observed allele frequency of allele *a* in pooled replicates of starting genotype *g* at site *s*

Using these values, we used the estimator of 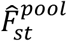defined in Hivert et al. 2018 to estimate *F*_*rg*_ for a single site:

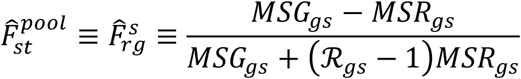

Where:

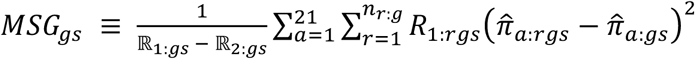 is the mean sum-of-squares for pooled replicates,

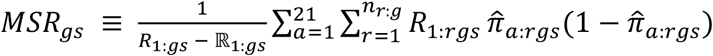 is the mean sum-of-squares within replicates,

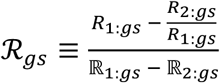, is the effective number of individuals after accounting for sampling, with 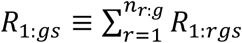,

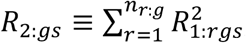,

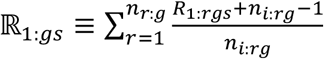, and

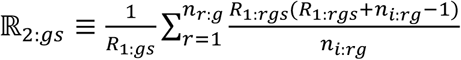.

Here, *n*_*r*:*g*_ is the number of replicates started from genotype *g* and *n*_*i*:*rg*_ is the number of individual phage in replicate *r* of starting genotype *g* in the sample used to make the sequencing library.

From the relationship between *C*_1_ and *F*_*rg*_ we approximated the site-specific effects of chance for a particular starting genotype as:

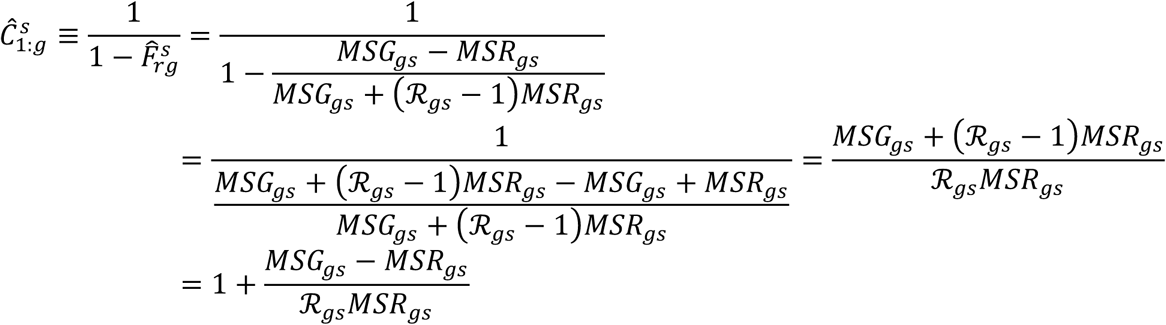

When there were replicates from more than one starting genotype, we calculated *MSG*_*gs*_, *MSR*_*gs*_, and *ℛ*_*gs*_ separately for each starting genotype and averaged these values together, using weights proportional to the number of replicates for that genotype. Thus:

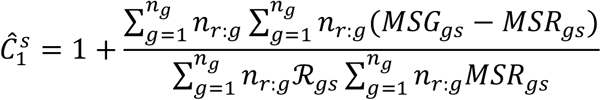

Where *n*_*g*_ is the number of distinct starting genotypes.

We then took the average numerator and average denominator as suggested by Hivert et al. 2018 (Hivert et al. 2018) and Weir and Cockerham 1984 for estimating *F*_*st*_:

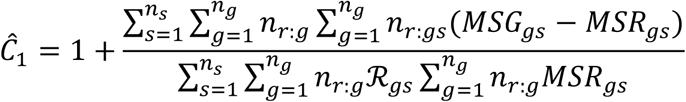

Where *n*_*s*_ is the number of sites.

#### Estimating the effects of contingency

To determine the effects of contingency (*C*_2_) on the outcomes of evolution, we compared the average allele frequency of replicate PACE experiments between different starting genotypes. For each starting genotype, we pooled all replicates started from that genotype and treated it as a single sample. We compared allele frequencies among genotypes by estimating the probability that two randomly chosen alleles in a sample would be different if they were both drawn from the same starting genotype (*V*_*g*_) or drawn from different starting genotypes (*V*_*t*_):

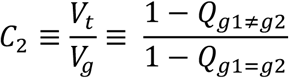

where *Q*_*g*1=*g*2_ is the probability that two randomly chosen alleles from the same starting genotype are identical in state and *Q*_*g*1≠*g*2_ is the probability that two randomly chosen alleles from different starting genotypes are identical in state. To calculate 1 − *Q*_*g*1≠*g*2_ we note that the probability of two randomly drawn alleles being different when chosen irrespective of starting genotype is simply the average of the probability of two randomly drawn alleles being different when they are drawn from the same and different starting genotypes, i.e.:

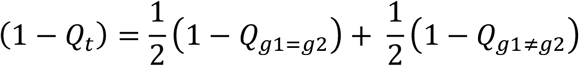

where *Q*_*t*_ is the probability that two randomly chosen alleles irrespective of starting genotype are identical in state. From this we have:

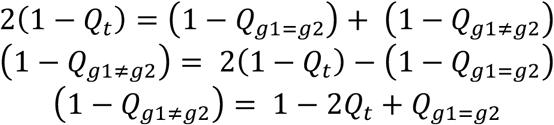

Using this and the fact that *Q*_*g*1=*g*2_ is equivalent to *Q*_*g*_ used above to calculate the effects of chance, we have:

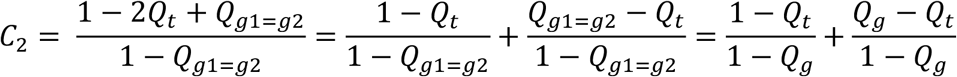

This statistic is related to Wright’s F_st_ as:

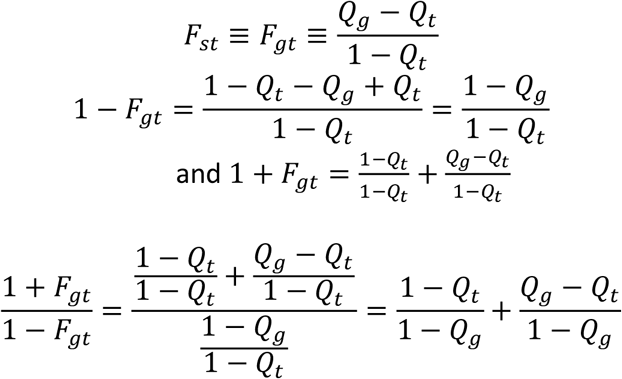

As with the effects of chance, we used the method of moments estimator defined in Hivert et al. 2018 to estimate the effects of contingency:

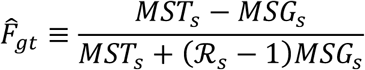

Where:

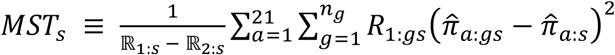 is the mean sum-of-squares for the entire pooled sample

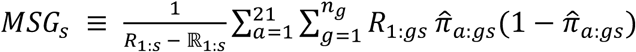 is the mean sum-of-squares for a starting genotype,

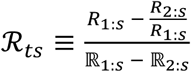, is the effective number of individuals after accounting for sampling, with 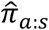 ≡ the observed allele frequency of allele *a* among all genotypes at site *s*,

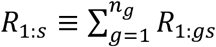,

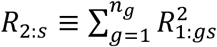,

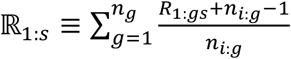, and

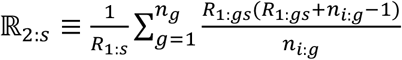.

With *n*_*i:g*_ being the number of individual phage used to make the libraries for starting genotype *g*.

From the relationship between *F*_*gt*_ and *C*_2_, we approximated the effects of contingency as:

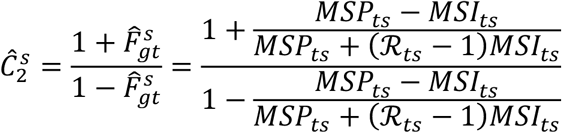

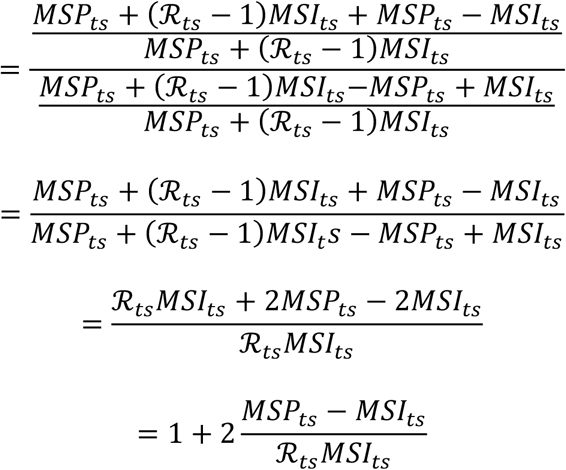

Again, we treated all sites as independent and summed the numerator and denominators to estimate the effects of contingency:

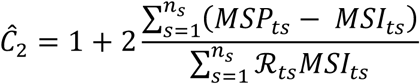

#### Estimating the combined effect of chance and contingency

To determine the combined effects of chance and contingency (*C*_3_) on the outcomes of evolution, we compared allele frequencies from individual replicates to the average allele frequency among replicates from different starting genotypes. In each case, we pooled replicates started from a genotype and treated it as a single sample and compared it to the individual replicates started from different genotypes. We compared allele frequencies by estimating the probability that two randomly chosen alleles would be different if they were both drawn from the same replicate or if they were drawn from a different starting genotype:

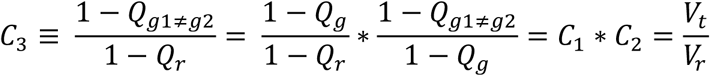

We thus used:

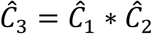

as our estimate of the combined effects of chance and contingency. This estimator indicates that the combined effects of chance and contingency are multiplicative and thus amplify each other’s effects as they get larger.

## Data and Code Availability

The high throughput sequencing data of evolved BCL-2 family protein variants were deposited in the National Center for Biotechnology Information (NCBI) Sequence Read Archive (SRA) databases. They can be accessed via BioProject: PRJNA647218. The processed sequencing data are available on Dryad (https://datadryad.org/stash/share/Ty32n-2c8nUZuiFegxDdy7wNeGDEsqlwtL8tOKz1RWU). The coding scripts and reference sequences for processing the data are available on Github (https://github.com/JoeThorntonLab/BCL2.ChanceAndContingency).

## ACKNOWLEDGEMENTS

We thank members of the Dickinson and Thornton groups for helpful comments on the manuscript, S. Ahmadiantehrani for editing, and R. Ranganathan for the use of the Illumina MiSeq instrument. This work was supported by CAREER Award 1749364 from the National Science Foundation (NSF) to B.C.D and National Institutes of Health grants R01GM131128 and R01GM121931 to J.W.T. V.C.X was supported by an NSF Graduate Research Fellowship (DGE-1746045). B.P.H.M. was supported by a NIH NRSA award (F32GM122251).

## AUTHOR CONTRIBUTIONS

All authors contributed to conception of the project. BCD, VCX, and JP designed the PACE dual-selection system, which VCX and JP engineered, optimized, and implemented. VCX and JP performed PACE, biochemical assays, and sequencing experiments, with input from BPHM. BPHM and JWT developed and designed the evolutionary and genetic analyses. BPHM led and performed the phylogenetic, genetic, and evolutionary analyses, with input from VCX and JP. All authors contributed to writing the manuscript.

---

**Figure S1.**
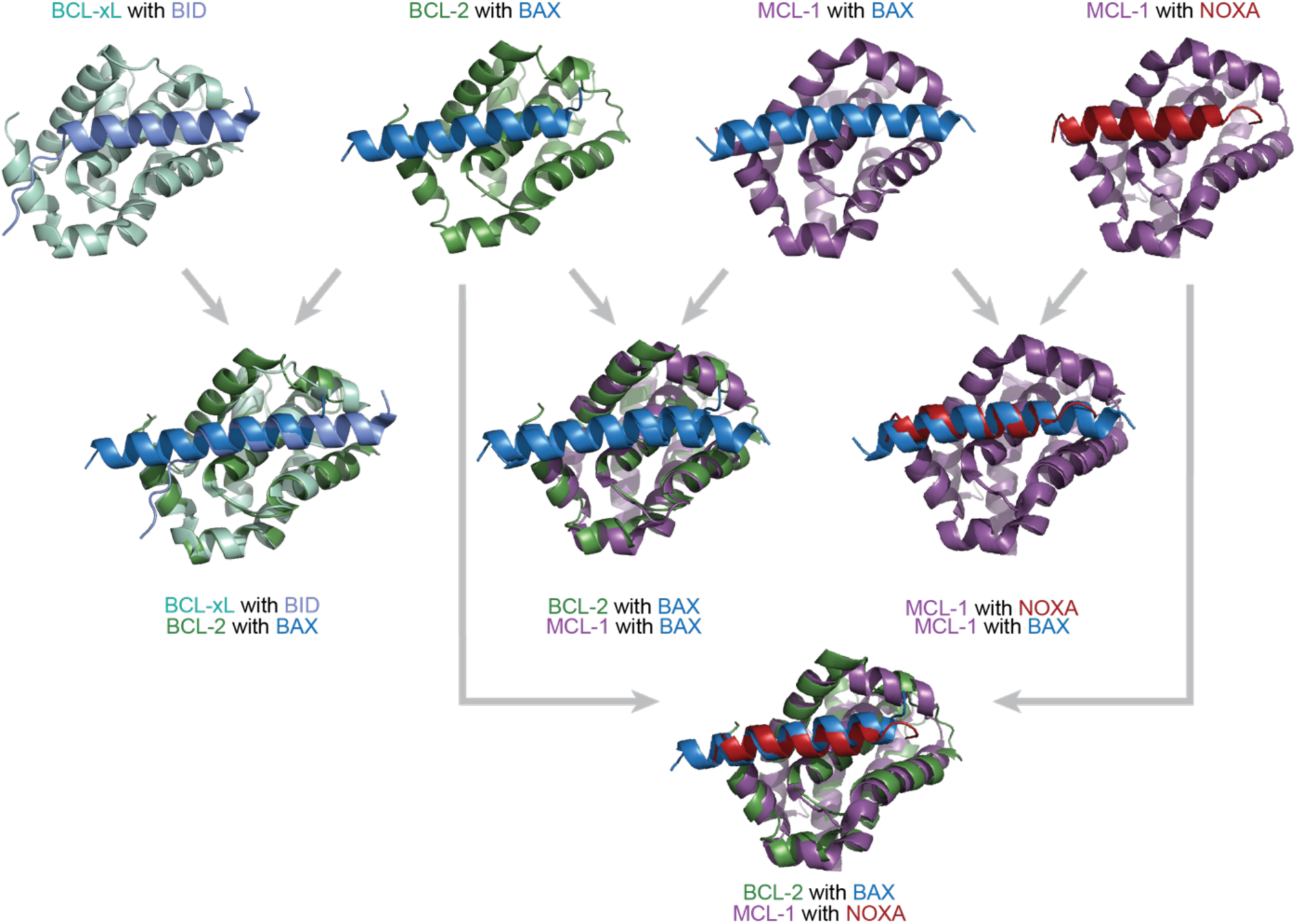
BCL-2 family proteins are structurally similar but have different binding profiles, Related to Figure 1. Crystal structures and overlays of BCL-xL (a vertebrate paralog of BCL-2, light green) bound to BID (light blue; PDB: 4qve); BCL-2 (green) bound to BAX (a protein with a BID-like binding profile, blue; PDB: 2xa0); MCL-1 (purple) bound to BAX (blue; PDB: 3pk1); and MCL-1 bound to NOXA (red; PDB: 2nla). The BCL-2 family proteins bind the coregulator proteins at the same interface.

**Figure S2.**
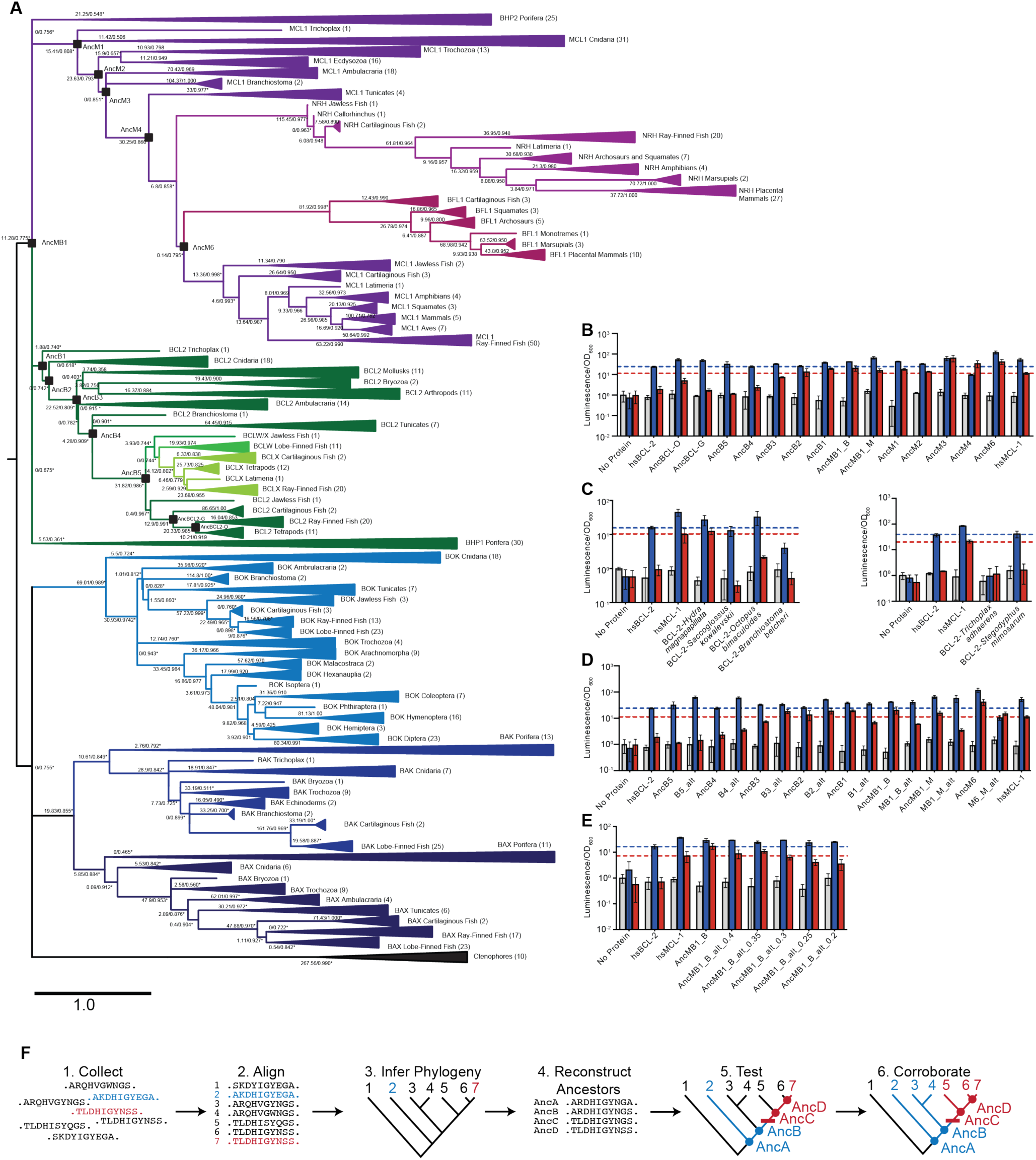
Phylogenetic reconstruction of the BCL-2 family proteins, Related to Figure 2. (A) Inferred phylogenetic relationships among BCL-2 family proteins. Green, BCL-2 class; purple, MCL-1 class; black, ctenophore sequences. Blue, pro-apoptotic paralogs. Shades within each color group indicate paralogs. Parentheses, number of sequences in each clade. Black squares, ancestral sequences reconstructed and tested. Node labels, approximate LRT statistics and transfer bootstrap values. Nodes with an asterisk were constrained to be congruent with known taxonomic relationships. (B) Interactions of ancestral reconstructed proteins with BID (blue) and NOXA (red) in the luciferase assay, compared to no-coregulator control (gray). Activity is scaled relative to no-coregulator control with no-BCL-2 family protein. Columns and error bars, mean ± SD of three biological replicates. hsBCL-2 with BID (dashed blue line). hsMCL-1 with (dashed red line). (C) Same as B, but for extant species *Hydra magnapapillata* (Cnidaria), *Octopus bimaculoides* (Lophotrochozoa), *Saccoglossus kowalevskii* (Hemichordata), *Branchiostoma belcheri* (Cephalochordata), *Trichoplax adhaerens* (Placozoa) and *Stegodyphus mimosarum* (Ecdysozoa). (D) Same as B, but contains alternative reconstructions (Alt) for each ancestor which combine all plausible alternative amino acid states (PP>0.2) in a single “worst-case” alternative reconstruction. (E) Same as B, but contains multiple alternative reconstructions for AncMB1_B. In each case, all plausible alternative amino acid states greater than the listed value are included in a single “worst-case” alternative reconstruction. (F) Ancestral state reconstruction. 1) Sequences conferring different functions (red v. blue) are collected, as well as related sequences whose function may be unknown (black). These sequences can be orthologs from a variety a species, paralogs from gene duplication events, or a combination of orthologs and paralogs. 2) Sequences are aligned to identify homologous sites and poorly aligned regions are removed. 3) A phylogeny is inferred. Additional sequences are added to fill in missing taxonomic groups and to help break apart long branches. If clear discrepancies between known species relationships and the inferred phylogeny remain, the phylogeny may be constrained to minimize the inferred number of gene duplication or loss events. The last common ancestor of sequences conferring different functions cannot be the root of the phylogeny. Instead, an outgroup is required (e.g. sequence 1) based on prior information. 4) Using the inferred phylogeny, the aligned sequences, and a model of sequence evolution, the most likely state at each ancestral node is determined. 5) Ancestral sequences between the sequences with known functional differences are synthesized and tested for function. 6) The function of ancestral proteins proposes an evolutionary hypothesis about the branch in which function was altered during history (red bar). Corroboration for this hypothesis can be gained by determining the function of extant sequences which bracket the specific branch in question (e.g. sequences 3, 4, 5, and 6).

**Figure S3.**
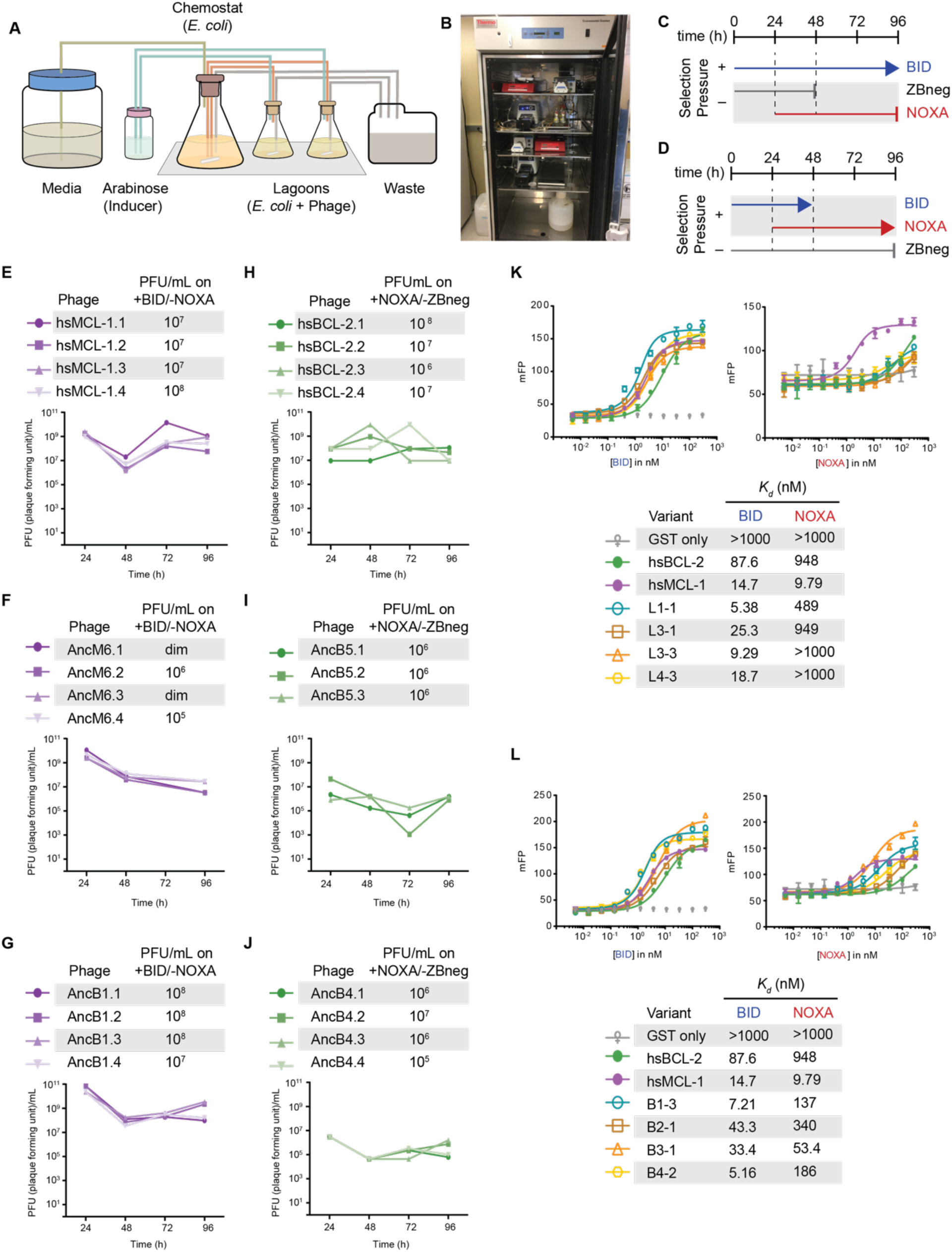
Using PACE to evolve target PPI binding specificity of BCL-2 family proteins, Related to Figure 3. (A) Schematic of a PACE experiment. Davis Rich carboy media flows into the chemostat, which contains *E. coli* with the positive selection (+AP), counter selection (-AP), and mutagenesis plasmids (MP). The cells then flow into the lagoons, which contain phage with the evolving BCL-2 family protein. Arabinose is pumped into the lagoons to induce the mutagenesis plasmid in the *E. coli*. Both chemostats and lagoons are connected to the waste to maintain proper volume, cell density, and flow rate. (B) Picture of representative PACE experiment from this work. (C) Timeline of PACE experiments where hsMCL-1, AncM6, and AncB1 were evolved to lose NOXA binding. ZBneg is a control zipper peptide. (D) Timeline of PACE experiments where hsBCL-2, AncB5, and AncB4 were evolved to gain NOXA binding. (E) Phage titers (PFU/mL) over time (bottom) and activity-dependent phage titers at the end of the PACE experiments (top) where hsMCL-1 was evolved to lose NOXA binding. Activity-dependent plaque assays used plasmids 28-46 and Jin 487. (F) Same as (E) for AncM6. “dim” means plaques were visible but weak, and therefore not quantifiable. (G) Same as (E) for AncB1. (H) Phage titers (PFU/mL) over time (bottom) and activity-dependent phage titers at the end of the PACE experiments (top) when hsBCL-2 was evolved to gain NOXA binding. Activity-dependent plaque assays used plasmids 28-48 and 29-39. (I) Same as (H) for AncB5. (J) Same as (H) for AncB4. (K) Fluorescence polarization of hsMCL-1 variants evolved to lose NOXA binding. Bars are the mean of three replicates; error bars, SD. mFP, normalized measured fluorescent polarization. (L) Fluorescence polarization of hsBCL-2 variants evolved to gain NOXA binding, same as (K).

**Figure S4.**
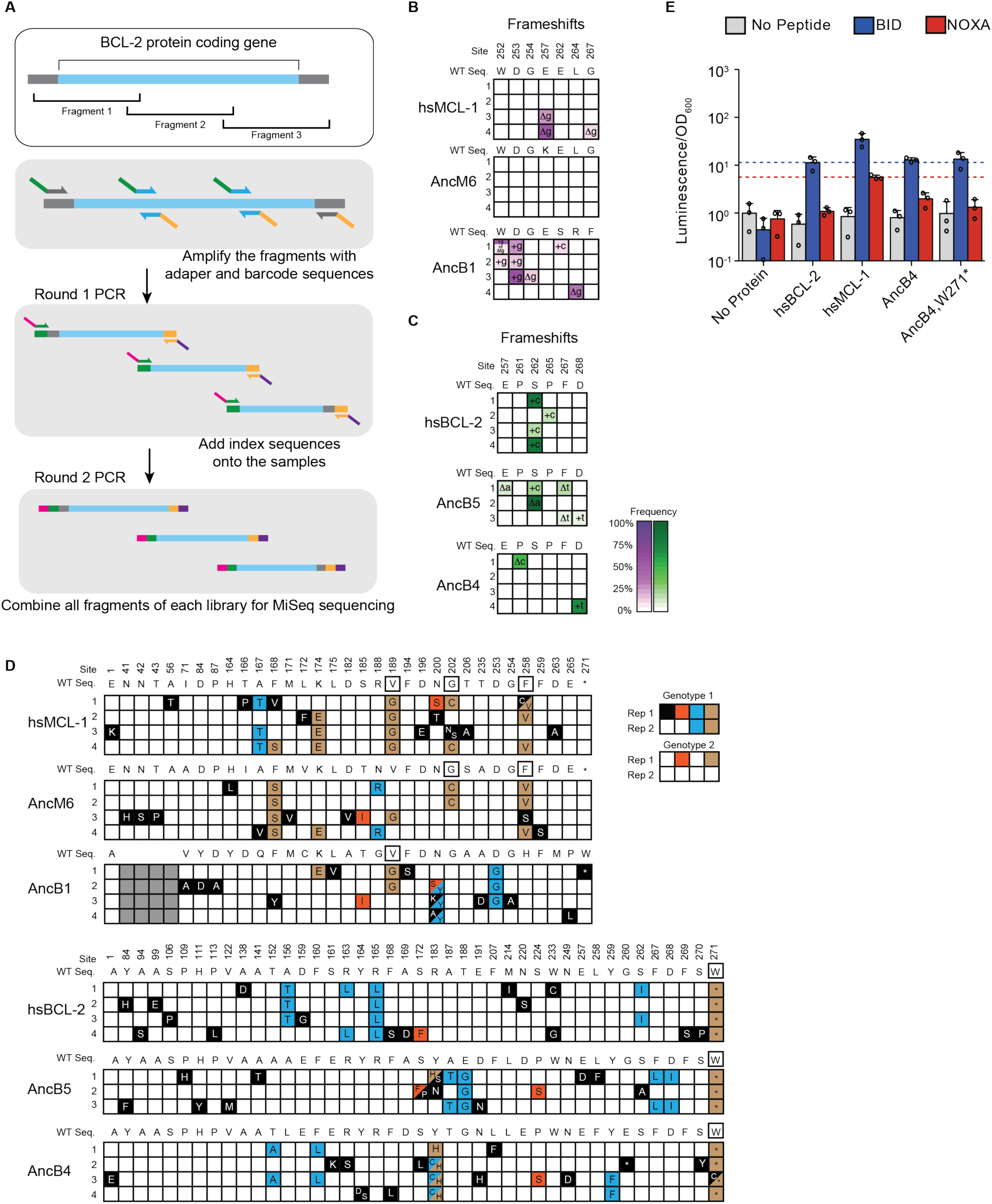
Allele frequency and mutational analysis of evolutionary outcomes, relate to Figure 4. (A) MiSeq library preparation. After isolation of phage DNA, the coding region of the evolving BCL-2 family protein was amplified in three overlapping fragments, each of which was smaller than 300bp. The DNA fragments were then amplified using sequence-specific primers. MiSeq sequence adapters were added in a second PCR step. These fragmented DNA libraries were combined and used for MiSeq high throughput sequencing. Blue, target gene coding region. Gray, adjacent vector sequence. Green, forward adapter and barcode sequence. Orange, reverse adapter and barcode sequence. Magenta, index 1 sequence. Purple, index 2 sequence. (B) Allele frequency data of frameshifts from replicate PACE experiments started from hsMCL-1, AncM6, and AncB1 evolved to lose NOXA binding. Site numbers and wildtype (WT) amino acid states are listed above each sequence. Each row represents an independent replicate population. Non-wildtype insertions and deletions that reached > 5% in frequency are shown, with frequency proportional to color saturation. Split cells show populations with multiple non-WT states > 5%. Plus (+) indicates an addition of a nucleotide. Delta (Δ) indicates a deletion of a nucleotide. (C) Same as (B) for replicate PACE experiments for hsBCL-2, AncB5, and AncB4 evolved to gain NOXA binding. (D) Categories of the 100 non-WT states observed for each non-WT state. Black box with white letters, mutant states observed in only one replicate. Teal, mutant states observed in multiple replicates from the same starting genotype. Orange, mutant states observed in a single replicate from multiple different starting genotypes. Brown, mutant states observed in multiple replicates from the same starting genotype and in at least one other replicate from a different starting genotype. Black box outline, mutant states observed in multiple replicates from the same starting genotype and from multiple replicates from a different starting genotype. Gray boxes are sites that do not exist in a particular sequence. (E) Luciferase assay of the w271∗ mutation in AncB4. Activity is scaled relative to the no BCL-2 family protein with no coregulator peptide control. Bars are the mean ± SD of three biological replicates (circles). Gray bar, no coregulator peptide. Blue bar, BID. Red bar, NOXA. Blue dotted lines mark the average signal of hsBCL-2 with BID, and red dotted lines mark the average signal of hsMCL-1 with NOXA.

**Figure S5.**
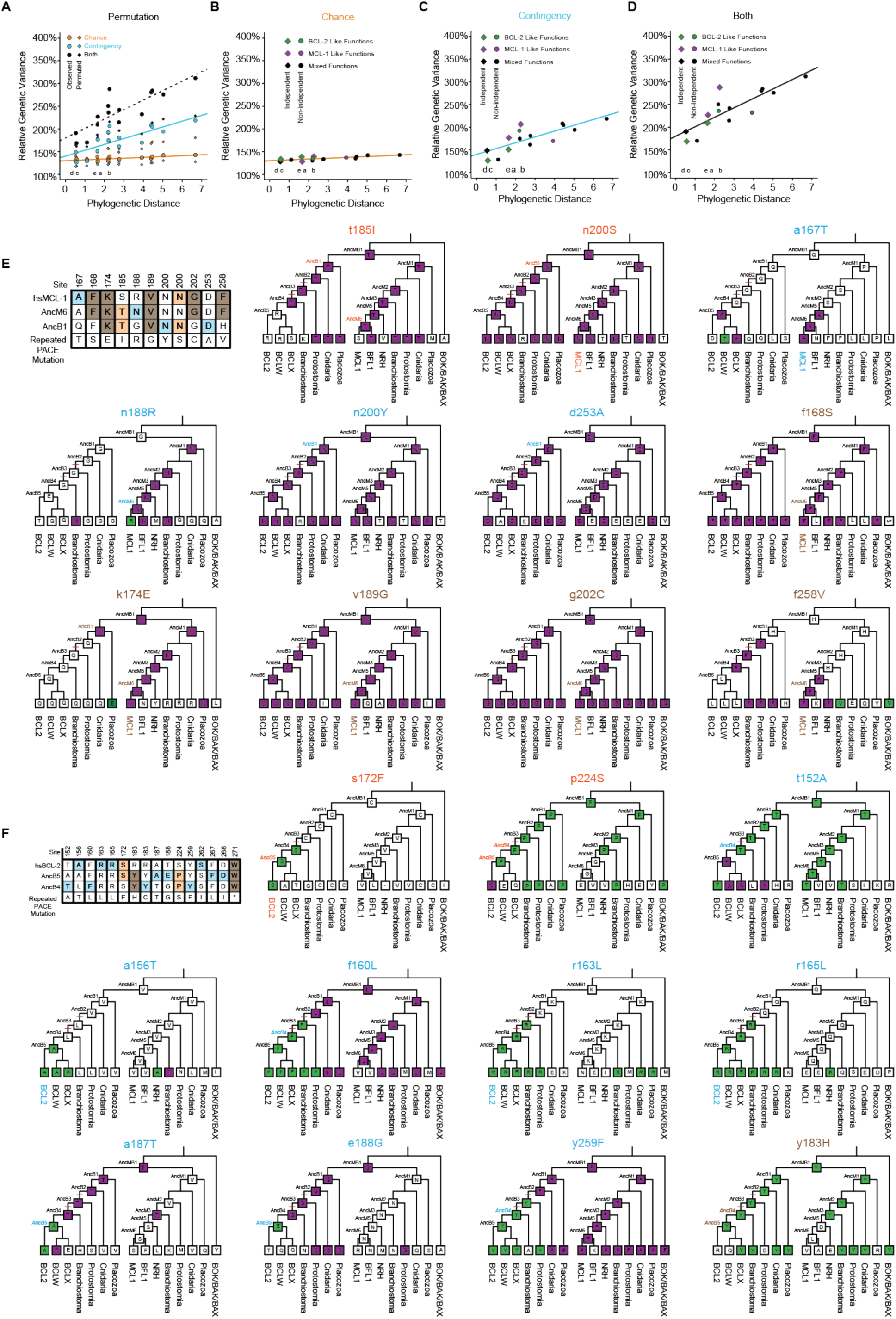
Chance and contingency in the repeated evolution of extant and ancestral BCL-2 family proteins, Related to Figure 5. (A) Change in chance and contingency over time. Relationship between phylogenetic distance between pairs of starting genotypes for experimental evolution (ancestral or extant proteins, as the total branch lengths separating them) and the effects of chance (orange), contingency (teal), or both (black) on the outcomes of evolution between them. Lines are best fits from linear models. Circles are observed values. Diamonds are averages of 1000 permutations of starting genotype labels. This shuffling of genotype labels results in more genetic variance among samples from the same ‘starting genotype’ than the observed data, and less genetic variance between samples from different ‘starting genotypes’ than the observed data. Letters indicate the specific branch from Figure 3D. (B) Change in chance over time. Green, both starting genotypes had BCL-2 like function. Purple, both starting genotypes had MCL-1 like function. Black, starting genotypes differed in function. Phylogenetically independent comparison are shown as diamonds. The effect of chance did not change with phylogenetic distance when restricting analysis to comparisons that are phylogenetically independent (slope=0.042, P=0.71) and genotypes selected for the same function (slope=0.029, P=0.82). (C) Change in contingency over time. Green, both starting genotypes had BCL-2 like function. Purple, both starting genotypes had MCL-1 like function. Black, starting genotypes differed in function. Phylogenetically independent comparison are shown as diamonds. The effect of contingency increased with phylogenetic distance and was marginally significant when restricting analysis to comparisons that are phylogenetically independent (slope=0.31, P=0.07), and genotypes selected for the same function (slope=0.42, P=0.05). (D) Change in the combined effect of chance and contingency over time. Green, both starting genotypes had BCL-2 like function. Purple, both starting genotypes had MCL-1 like function. Black, starting genotypes differed in function. Phylogenetically independent comparison are shown as diamonds. The combined effect of chance and contingency increased with phylogenetic distance when restricting analysis to comparisons that are phylogenetically independent (slope=0.50, P=0.009) and genotypes selected for the same function (slope=0.63, P=0.01). (E) Phylogenetic distribution of repeated PACE mutations when selecting hsMCL-1, AncM6, and AncB1 against NOXA binding. Repeated mutation state and position are given above each cladogram. Lowercase letters, WT state. Uppercase letters, repeated mutant state. Top left: WT states for hsMCL-1, AncM6, and AncB1 at sites where repeated PACE mutations occurred. Repeated PACE mutation state is given in the last row. WT states are colored based on the pattern of repeated PACE mutations. Teal, sites that evolved the same mutation in multiple replicates from the same starting genotype. Orange, sites that evolved the same mutation in a single replicate from multiple starting genotypes. Brown, sites that evolved the same mutation in multiple replicates from the same starting genotype and from multiple different starting genotypes. Starting genotypes for PACE are colored similarly. Each cladogram shows the WT state for each node. Purple boxes; same WT state as the sequence in which the repeated PACE mutation emerged. Green boxes; same WT state as the repeated PACE mutation. Red bars indicate the interval in which NOXA binding was historically lost. (F) Same as (E) when selecting hsBCL-2, AncB5, and AncB4 for NOXA binding. Green boxes; same WT state as the sequence in which the repeated PACE mutation emerged. Purple boxes; same WT state as the repeated PACE mutation.

**Figure S6.**
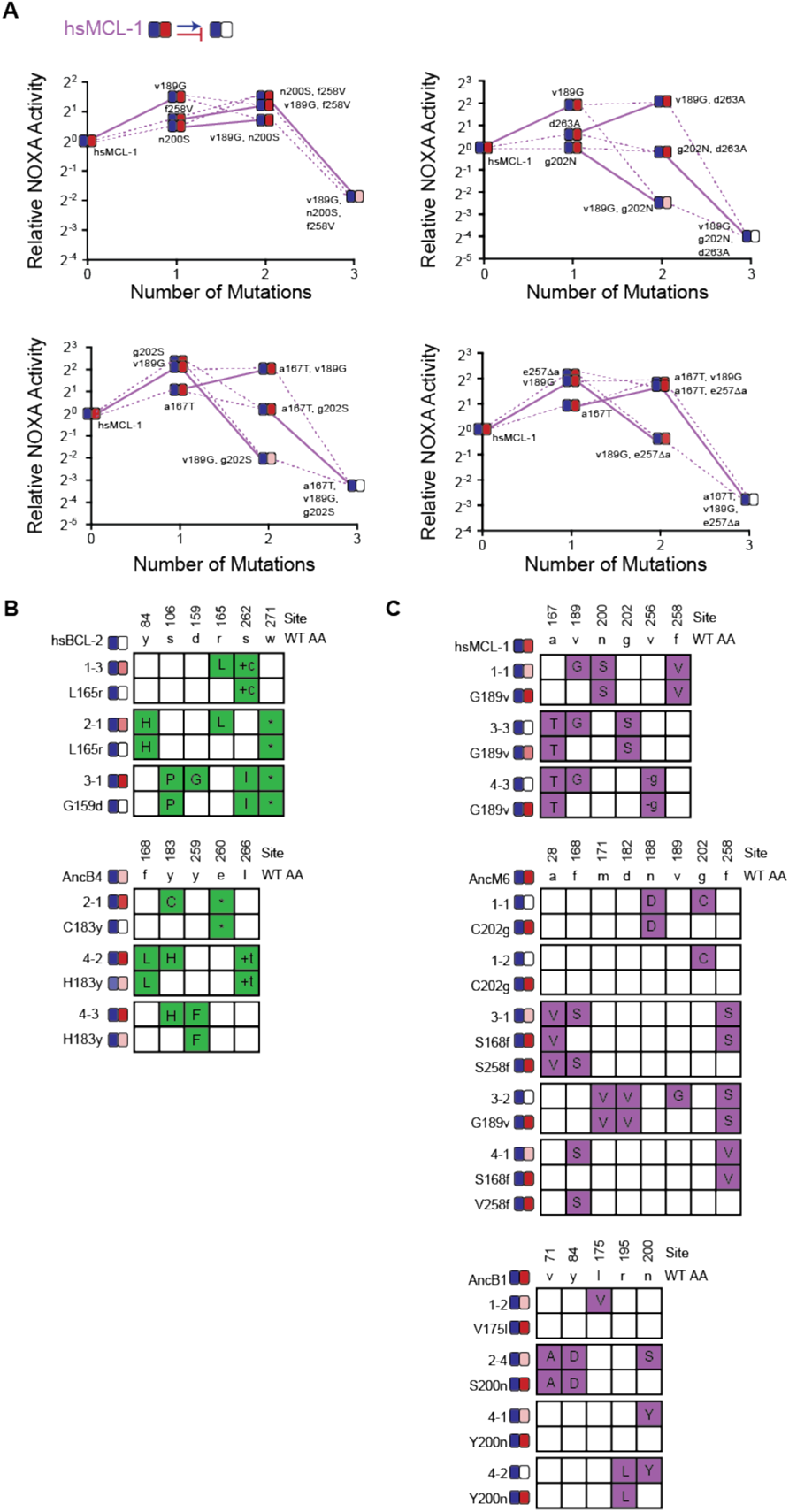
Effects of individual PACE-derived mutations, Related to Figure 6. (A) Effect on NOXA binding of mutations that occurred in PACE variants when hsMCL-1 was evolved to lose NOXA binding. Each panel shows NOXA binding (y-axis) for a unique variant as additional mutations are added (x-axis). Values are the mean of three biological replicates. Heatmaps show the effects of each mutation on BID (blue) and NOXA (red) activity, and each shaded box represents the normalized mean of three biological replicates. Lines connect genotypes that differ by a single mutation. Solid lines show the effects of the v189G mutation. Dashed lines show the effects of all other mutations. Mutations come from variants L1-1 (top left), L3-1 (top right), L3-3 (bottom left), and L4-3 (bottom right). (B) Phenotypic effects of reverting frequent PACE-derived mutations. Individual variants were isolated from PACE experiments that selected for the gain of NOXA binding in hsBCL-2 and AncB4. Sites and WT amino state are indicated at top. For each variant, non-WT states are shown in green. Heatmaps on the left show binding to BID and NOXA in the luciferase assay for each variant and their corresponding mutant without the key mutation. Each shaded box represents the normalized mean of three biological replicates. (C) Same as (B), but for the loss of NOXA binding in hsMCL-1, AncM6, and AncB1. For each variant, non-WT states are shown in purple.

**Figure S7.**
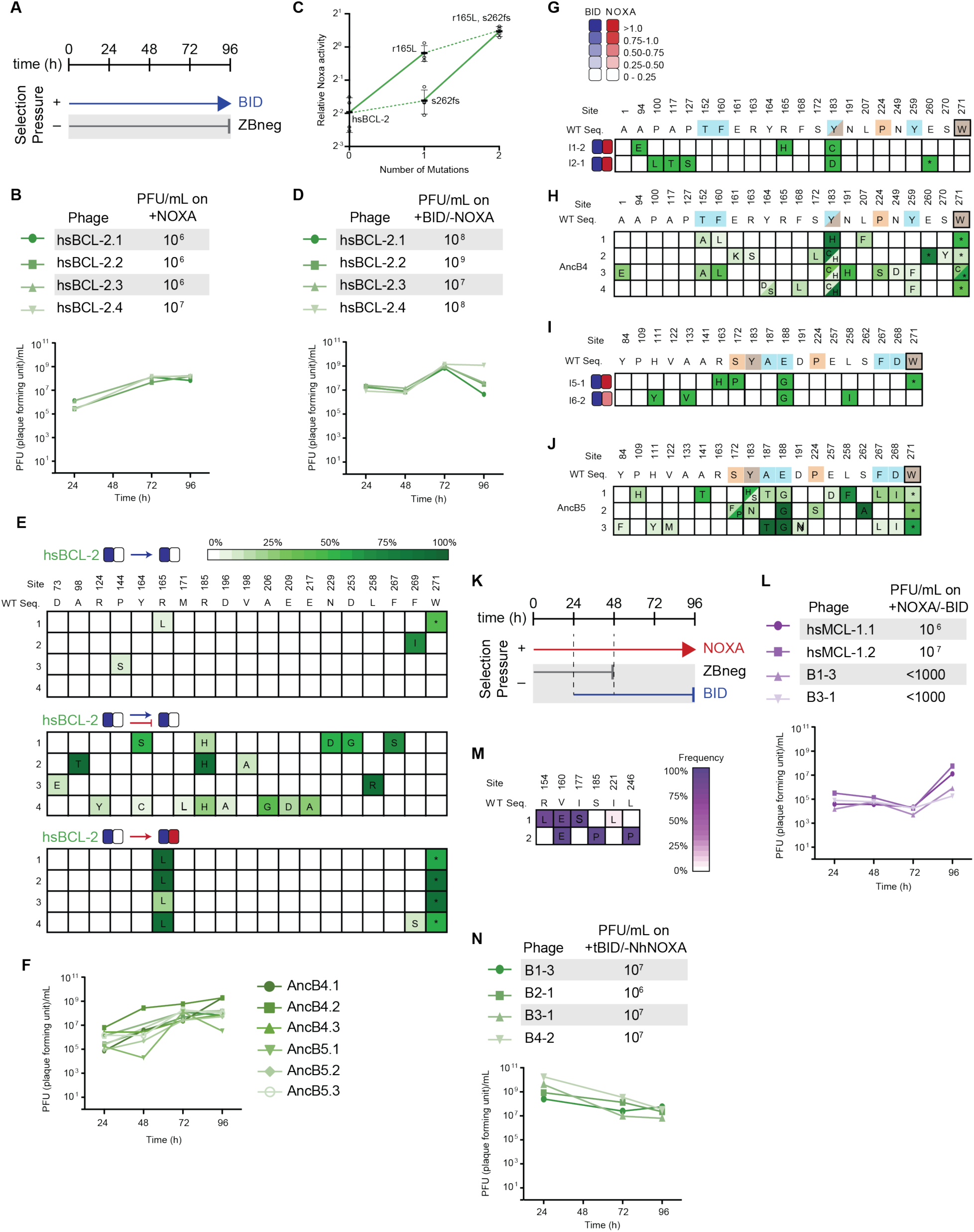
Spontaneous emergence of NOXA binding and selection for NOXA-specific binders, Related to Figure 7. (A) Timeline of PACE experiments where hsBCL-2, AncB5, and AncB4 were evolved with only positive selection to maintain BID binding. Selection conditions shown as arrows and blunt bars: arrow, selection for binding to BID (blue); blunt bar, selection against binding to ZBneg (gray). (B) Phage titers (PFU/mL) over time (bottom) and activity-dependent phage titers at the end of the PACE experiment (top) where hsBCL-2 was evolved to maintain BID binding. Activity-dependent plaque assays used plasmid 28-48. (C) Effect on NOXA binding of the key r165L mutation. Bars are the mean ± SD of three biological replicates (circles). Solid lines show the effects of the r165L mutation while dotted lines show the effect of a frameshift (fs) at site 262. (D) Same as (B) where hsBCL-2 was evolved for binding and against NOXA binding. Activity-dependent plaque assays used plasmids 28-48 and Jin 487. (E) Allele frequency of non-wildtype states when hsBCL-2 was evolved to maintain BID binding (top) or when hsBCL-2 was evolved to simultaneously maintain BID binding and lose NOXA binding (middle). For comparison, the same sites are also shown for when hsBCL-2 was evolved to gain NOXA binding (bottom). Site numbers and wildtype (WT) amino acid states are listed above each sequence. Each row represents an independent replicate population. Non-wildtype amino acids that reached > 5% in frequency are shown, with frequency proportional to color saturation. (F) Phage titers (PFU/mL) over time from the PACE experiment where AncB4 and AncB5 were evolved to maintain BID binding. (G) Phenotypes and genotypes of individual AncB4 variants that were isolated from PACE when selecting for BID binding and screened for the gain of NOXA binding. Site numbers and wildtype (WT) amino acid states are indicated at the top. Heatmaps on the left show binding to BID (blue) and NOXA (red) in the luciferase assay for each variant, and each shaded box represents the normalized mean of three biological replicates. (H) Non-wildtype amino acid frameshifts that reached > 5% in frequency are shown for PACE where AncB4 was evolved to gain NOXA binding, for comparison with (F). Frequency is proportional to color saturation. Split cells show populations with multiple non-WT states > 5%. Each row represents an independent replicate lagoon. Color of WT state indicate if the mutation was seen among multiple replicates of the same starting genotype (teal), a single replicate from multiple starting genotypes (orange), or in multiple replicates and multiple starting genotypes (brown). Black box outline indicates mutant states observed in multiple replicates from the same starting genotype and from multiple replicates from a different starting genotype (I) Same as (F) but for AncB5. (J) Same as (G) but for AncB5 and for comparison with (I). (K) Timeline of PACE experiments where hsMCL-1 and two previously-evolved NOXA-binding hsBCL-2 variants were evolved to maintain NOXA binding and lose BID binding. Selection conditions: arrow, selection for binding NOXA (red); blunt bar, selection against binding a specific peptide (BID (blue) or ZBneg (gray)). (L) Phage titers (PFU/mL) over time (bottom) and activity-dependent phage titers at the end of the PACE experiment (top) where hsMCL-1 and NOXA-binding hsBCL-2 variants were evolved for binding NOXA and against BID. Activity-dependent plaque assays used plasmids 28-48 and Jin 518. Limit of detection = 10^3^ PFU/mL. (M) Allele frequency of non-wildtype states after hsMCL-1 was evolved to maintain NOXA binding and lose BID binding. Site numbers and wildtype (WT) amino acid states are listed above each sequence. Each row represents an independent replicate lagoon. Non-wildtype amino acid frameshifts that reached > 5% in frequency are shown, with frequency proportional to color saturation. (N) Phage titers (PFU/mL) over time (bottom) and activity-dependent phage titers at the end of the PACE experiment (top) where NOXA-binding hsBCL-2 variants were evolved to lose NOXA binding. Activity-dependent plaque assays used plasmids 28-46 and Jin 487.

**Table S1 (Excel File)**. Luciferase assay data for all experiments. Related to Figures 1, 2, 3, 6, 7, S4, S6, S7.

**Table S2 (Excel File)**. Posterior probabilities for reconstructed ancestral sequences. Related to Figure 2. For each sequence, the site, maximum likelihood (ML) amino acid state, and posterior probability (PP) are given, along with the highest posterior probability alternative (ALT) state and posterior probability for this alternative state. Locations of paralog specific insertions are shown as gaps. For each reconstructed sequence, the average posterior probability for the maximum likelihood states and the alternative states is given, as are the number of sites where the posterior probability of a non-maximum likelihood state is greater than 0.2. Finally, the average, maximum, minimum, and variance among reconstructed ancestors is given for the average maximum likelihood posterior probability and the number of non-maximum likelihood states greater than 0.2 posterior probability.

**Table S3 (Excel File)**. List of PACE experiments, amino acid alignments of hsBCL-2 and hsMCL-1 with their structural global alignment, and mutations found in individual variants isolated from PACE. fs is frameshift, aa is amino acid, co is codon change. Related to STAR Methods.

**Table S4 (Excel file)**. PACE library and high-throughput sequencing (HTS) data. Related to STAR Methods. PACE experiments are listed in the tab “Library-info” which contains the name, purpose of the experiment, and HTS experiment numbers. The tab “Primers for HTS” lists all the primer sequences used for HTS library constructions. The tab “MiSeq reads number” include the read number of each library in this MiSeq run and the library sample information. The library samples are labeled as X∗-end or X∗-$$. “X” indicates the specific PACE experiment, “∗” the experimental replicate, “end” means samples were collected after 96 hours when the experiment finished, and “$$” indicates the time point after removing chemostat A (e.g., “B2-24” is a sample from replicate 2 of evolution B and collected 24 hours after removing chemostat A, which is 72 hours from the start of PACE). The tab “genotype” includes the aligned protein sequences with corresponding residue numbers. The ‘Frequency’ tab contains the non-wildtype amino acid frequency of each sample for each site.

**Table S5 (Excel file)**. Descriptions of plasmids and sequences used. Related to STAR Methods.

## REFERENCES

Altschul, S. F., T. L. Madden, A. A. Schaffer, J. Zhang, Z. Zhang, W. Miller, and D. Lipman. 1997. Gapped BLAST and PSI-BLAST: a new generation of protein database search programs. Nucleic Acids Res. 25:3389–3402.

Arendt, J., and D. Reznick. 2008. Convergence and parallelism reconsidered: what have we learned about the genetics of adaptation? Trends Ecol. Evol. 23:26–32.

Badran, A. H., V. M. Guzov, Q. Huai, M. M. Kemp, P. Vishwanath, W. Kain, A. M. Nance, A. Evdokimov, F. Moshiri, K. H. Turner, P. Wang, T. Malvar, and D. R. Liu. 2016. Continuous evolution of Bacillus thuringiensis toxins overcomes insect resistance. Nature 533:58–63. Nature Publishing Group.

Badran, A. H., and D. R. Liu. 2015. Development of potent in vivo mutagenesis plasmids with broad mutational spectra. Nat. Commun. 6:8425. Nature Publishing Group.

Baier, F., N. Hong, G. Yang, A. Pabis, A. Barrozo, and D. Paul. 2019. Cryptic genetic variation shapes the adaptive evolutionary potential of enzymes. Elife 8:1–20.

Banjara, S., C. D. Suraweera, M. G. Hinds, and M. Kvansakul. 2020. The Bcl-2 family: Ancient origins, conserved structures, and divergent mechanisms. Biomolecules 10:1–21.

Beatty, J. 2009. Chance Variation and Evolutionary Contingency: Darwin, Simpson, The Simpsons, and Gould. Oxford Handb. Philos. Biol. 1–22.

Bloom, J. D., L. I. Gong, and D. Baltimore. 2010. Permissive secondary mutations enable the evolution of influenza oseltamivir resistance. Science 328:1272–1275.

Blount, Z. D., J. E. Barrick, C. J. Davidson, and R. E. Lenski. 2012. Genomic analysis of a key innovation in an experimental Escherichia coli population. Nature 489:513–518. Nature Publishing Group.

Blount, Z. D., C. Z. Borland, and R. E. Lenski. 2008. Historical contingency and the evolution of a key innovation in an experimental population of Escherichia coli. Proc. Natl. Acad. Sci. 105:7899–7906.

Blount, Z. D., R. E. Lenski, and J. B. Losos. 2018. Contingency and determinism in evolution: Replaying life’s tape. Science 362:eaam5979.

Bollback, J. P., and J. P. Huelsenbeck. 2009. Parallel genetic evolution within and between bacteriophage species of varying degrees of divergence. Genetics 181:225–234.

Breen, M. S., C. Kemena, P. K. Vlasov, C. Notredame, and F. A. Kondrashov. 2012. Epistasis as the primary factor in molecular evolution. Nature 490:535–538. Nature Publishing Group.

Bridgham, J. T., E. a Ortlund, and J. W. Thornton. 2009. An epistatic ratchet constrains the direction of glucocorticoid receptor evolution. Nature 461:515–519. Nature Publishing Group.

Carlson, J. C., A. H. Badran, D. A. Guggiana-Nilo, and D. R. Liu. 2014. Negative selection and stringency modulation in phage-assisted continuous evolution. Nat. Chem. Biol. 10:216–222. Nature Publishing Group, a division of Macmillan Publishers Limited. All Rights Reserved.

Certo, M., V. D. G. Moore, M. Nishino, G. Wei, S. Korsmeyer, S. A. Armstrong, and A. Letai. 2006. Mitochondria primed by death signals determine cellular addiction to antiapoptotic BCL-2 family members. Cancer Cell 9:351–365.

Chandler, C. H., S. Chari, and I. Dworkin. 2013. Does your gene need a background check? How genetic background impacts the analysis of mutations, genes, and evolution. Trends Genet. 29:358–366. Elsevier Ltd.

Chen, L., S. N. Willis, A. Wei, B. J. Smith, J. I. Fletcher, M. G. Hinds, P. M. Colman, C. L. Day, J. M. Adams, and D. C. S. Huang. 2005. Differential targeting of prosurvival Bcl-2 proteins by their BH3-only ligands allows complementary apoptotic function. Mol. Cell 17:393–403.

Chipuk, J. E., T. Moldoveanu, F. Llambi, M. J. Parsons, and D. R. Green. 2010. The BCL-2 Family Reunion. Mol. Cell 37:299–310. Elsevier Inc.

Couñago, R., S. Chen, and Y. Shamoo. 2006. In Vivo Molecular Evolution Reveals Biophysical Origins of Organismal Fitness. Mol. Cell 22:441–449.

Danial, N. N., and S. J. Korsmeyer. 2004. Cell Death: Critical Control Points. Cell 116:205–219.

De Laval, V. R., G. Deléage, A. Aouacheria, and C. Combet. 2014. BCL2DB: Database of BCL-2 family members and BH3-only proteins. Database 2014:1–7.

Delsuc, F., H. Philippe, G. Tsagkogeorga, P. Simion, M. K. Tilak, X. Turon, S. López-Legentil, J. Piette, P. Lemaire, and E. J. P. Douzery. 2018. A phylogenomic framework and timescale for comparative studies of tunicates. BMC Biol. 16:1–14. BMC Biology.

Dickinson, B. C., A. M. Leconte, B. Allen, K. M. Esvelt, and D. R. Liu. 2013. Experimental interrogation of the path dependence and stochasticity of protein evolution using phage-assisted continuous evolution. Proc. Natl. Acad. Sci. 110:9007–9012.

Dutta, S., S. Gullá, T. S. Chen, E. Fire, R. A. Grant, and A. E. Keating. 2010. Determinants of BH3 Binding Specificity for Mcl-1 versus Bcl-xL. J. Mol. Biol. 398:747–762. Elsevier Ltd.

Echave, J., S. J. Spielman, and C. O. Wilke. 2016. Causes of evolutionary rate variation among protein sites. Nat. Rev. Genet. 17:109–121.

Esvelt, K. M., J. C. Carlson, and D. R. Liu. 2011. A system for the continuous directed evolution of biomolecules. Nature 472:499–503.

Finnigan, G. C., V. Hanson-Smith, T. H. Stevens, and J. W. Thornton. 2012. Evolution of increased complexity in a molecular machine. Nature 481:394–398. Nature Publishing Group.

Gompel, N., and B. Prud’homme. 2009. The causes of repeated genetic evolution. Dev. Biol. 332:36–47. Elsevier Inc.

Gompel, N., B. Prud’homme, P. J. Wittkopp, V. A. Kassner, and S. B. Carroll. 2005. Chance caught on the wing: cis-regulatory evolution and the origin of pigment patterns in Drosophila. Nature 433:481–487.

Gong, L. I., M. A. Suchard, and J. D. Bloom. 2013. Stability-mediated epistasis constrains the evolution of an influenza protein. Elife 2013:1–19.

Goodsell, D. S., and A. J. Olson. 2000. Structural Symmetry and Protein Function. Annu. Rev. Biophys. Biomol. Struct. 29:105–153.

Gould, S. J. 1990. Wonderful life: the Burgess Shale and the nature of history. Norton.

Gould, S. J., and R. C. Lewontin. 1979. The spandrels of San Marco and the Panglossian paradigm: a critique of the adaptationist programme. Proc. R. Soc. London. Ser. B, Biol. Sci. 205:581–98.

Harms, M. J., and J. W. Thornton. 2014. Historical contingency and its biophysical basis in glucocorticoid receptor evolution. Nature 512:203–207. Nature Publishing Group.

Hawkins, N. J., C. Bass, A. Dixon, and P. Neve. 2019. The evolutionary origins of pesticide resistance. Biol. Rev. 94:135–155.

Hivert, V., R. Leblois, E. J. Petit, M. Gautier, and R. Vitalis. 2018. Measuring Genetic Differentiation from Pool-seq Data. Genetics 210:genetics.300900.2018.

Hubbard, B. P., A. H. Badran, J. A. Zuris, J. P. Guilinger, K. M. Davis, L. Chen, S. Q. Tsai, J. D. Sander, J. K. Joung, and D. R. Liu. 2015. Continuous directed evolution of DNA-binding proteins to improve TALEN specificity. Nat. Methods 12:939–942. Nature Publishing Group, a division of Macmillan Publishers Limited. All Rights Reserved.

Hughes, L. C., G. Ortí, Y. Huang, Y. Sun, C. C. Baldwin, A. W. Thompson, D. Arcila, R. Betancur-R., C. Li, L. Becker, N. Bellora, X. Zhao, X. Li, M. Wang, C. Fang, B. Xie, Z. Zhou, H. Huang, S. Chen, B. Venkatesh, and Q. Shi. 2018. Comprehensive phylogeny of ray-finned fishes (Actinopterygii) based on transcriptomic and genomic data. Proc. Natl. Acad. Sci. 201719358.

Jablonski, D. 2017. Approaches to Macroevolution: 1. General Concepts and Origin of Variation. Evol. Biol. 44:427–450. Springer US.

Jensen, J. D., B. A. Payseur, W. Stephan, C. F. Aquadro, M. Lynch, D. Charlesworth, and B. Charlesworth. 2019. The importance of the Neutral Theory in 1968 and 50 years on: A response to Kern and Hahn 2018. Evolution 73:111–114.

Kacar, B., X. Ge, S. Sanyal, and E. A. Gaucher. 2017. Experimental Evolution of Escherichia coli Harboring an Ancient Translation Protein. J. Mol. Evol. 84:69–84. Springer US.

Kale, J., E. J. Osterlund, and D. W. Andrews. 2018. BCL-2 family proteins: Changing partners in the dance towards death. Cell Death Differ. 25:65–80. Nature Publishing Group.

Karageorgi, M., S. C. Groen, F. Sumbul, J. N. Pelaez, K. I. Verster, J. M. Aguilar, S. L. Bernstein, T. Matsunaga, M. Astourian, G. Guerra, F. Rico, S. Dobler, A. A. Agrawal, and N. K. Whiteman. 2019. Genome editing retraces the evolution of toxin resistance in the monarch butterfly. Nature 574:409–412.

Kern, A. D., and M. W. Hahn. 2018. The neutral theory in light of natural selection. Mol. Biol. Evol. 35:1366–1371.

Kimura, M. 1986. DNA and the Neutral Theory. Philos. Trans. R. Soc. B Biol. Sci. 312:343–354.

Kimura, M. 1983. The neutral theory of molecular evolution. Cambridge University Press.

Kimura, M., and T. Ota. 1974. On some principles governing molecular evolution. Proc. Natl. Acad. Sci. U. S. A. 71:2848–2852.

Kozlov, A. M., and A. Stamatakis. 2019. Using RAxML-NG in Practice. Preprints 1–25.

Kryazhimskiy, S., D. P. Rice, E. R. Jerison, and M. M. Desai. 2014. Global epistasis makes adaptation predictable despite sequence-level stochasticity. Science 344:1519–1522.

Lanave, C., M. Santamaria, and C. Saccone. 2004. Comparative genomics: The evolutionary history of the Bcl-2 family. Gene 333:71–79.

Lobkovsky, A. E., and E. V. Koonin. 2012. Replaying the tape of life: Quantification of the predictability of evolution. Front. Genet. 3:1–8.

Lomonosova, E., and G. Chinnadurai. 2008. BH3-only proteins in apoptosis and beyond: An overview. Oncogene 27:S2–S19. Nature Publishing Group.

Mayr, E. 1983. How to carry out the adaptationist program? Am. Nat. 121:324–334.

McKeown, A. N., J. T. Bridgham, D. W. Anderson, M. N. Murphy, E. A. Ortlund, and J. W. Thornton. 2014. Evolution of DNA Specificity in a Transcription Factor Family Produced a New Gene Regulatory Module. Cell 159:58–68. Elsevier Inc.

Menéndez-Arias, L. 2010. Molecular basis of human immunodeficiency virus drug resistance: An update. Antiviral Res. 85:210–231.

Meyer, J. R., D. T. Dobias, J. S. Weitz, J. E. Barrick, R. T. Quick, and R. E. Lenski. 2012. Repeatability and Contingency in the Evolution of a Key Innovation in Phage Lambda.

Miller, H. C., P. J. Biggs, C. Voelckel, and N. J. Nelson. 2012. De novo sequence assembly and characterisation of a partial transcriptome for an evolutionarily distinct reptile, the tuatara (Sphenodon punctatus). BMC Genomics 13.

Monod, J. 1972. Chance & Necessity. Translation of le hasard et la necessite. First Vintage Books Edition.

Moroz, L. L., K. M. Kocot, M. R. Citarella, S. Dosung, T. P. Norekian, I. S. Povolotskaya, A. P. Grigorenko, C. Dailey, E. Berezikov, K. M. Buckley, A. Ptitsyn, D. Reshetov, K. Mukherjee, T. P. Moroz, Y. Bobkova, F. Yu, V. V. Kapitonov, J. Jurka, Y. V. Bobkov, J. J. Swore, D. O. Girardo, A. Fodor, F. Gusev, R. Sanford, R. Bruders, E. Kittler, C. E. Mills, J. P. Rast, R. Derelle, V. V. Solovyev, F. A. Kondrashov, B. J. Swalla, J. V. Sweedler, E. I. Rogaev, K. M. Halanych, and A. B. Kohn. 2014. The ctenophore genome and the evolutionary origins of neural systems. Nature 510:109–114. Nature Publishing Group.

Natarajan, C., F. G. Hoffmann, R. E. Weber, A. Fago, C. C. Witt, and J. F. Storz. 2016. Predictable convergence in hemoglobin function has unpredictable molecular underpinnings. Science.

Nguyen, V., C. Wilson, M. Hoemberger, J. B. Stiller, R. V Agafonov, S. Kutter, J. English, D. L. Theobald, and D. Kern. 2016. Evolutionary drivers of thermoadaptation in enzyme catalysis. Science 3717:289–294.

Orgogozo, V. 2015. Replaying the tape of life in the twenty-first century. Interface Focus 5:20150057.

Ortlund, E. A., J. T. Bridgham, M. R. Redinbo, and J. W. Thornton. 2007. Crystal structure of an ancient protein: evolution by conformational epistasis. Science 317:1544–1548.

Perutz, M. F., J. C. Kendrew, and H. C. Watson. 1965. Structure and function of haemoglobin: II. Some relations between polypeptide chain configuration and amino acid sequence. J. Mol. Biol. 13:669–678. Academic Press Inc. (London) Ltd.

Petros, A. M., E. T. Olejniczak, and S. W. Fesik. 2004. Structural biology of the Bcl-2 family of proteins. Biochim. Biophys. Acta - Mol. Cell Res. 1644:83–94.

Pollock, D. D., G. Thiltgen, and R. A. Goldstein. 2012. Amino acid coevolution induces an evolutionary Stokes shift. Proc. Natl. Acad. Sci. U. S. A. 109.

Pu, J., J. A. Dewey, A. Hadji, J. L. Labelle, and B. C. Dickinson. 2017a. RNA Polymerase Tags to Monitor Multidimensional Protein-Protein Interactions Reveal Pharmacological Engagement of Bcl-2 Proteins. J. Am. Chem. Soc. 139:11964–11972.

Pu, J., M. Disare, and B. C. Dickinson. 2019. Evolution of C-Terminal Modification Tolerance in Full-Length and Split T7 RNA Polymerase Biosensors. ChemBioChem 20:1547–1553.

Pu, J., J. Zinkus-Boltz, B. C. Dickinson, J. Z.- Boltz, and B. C. Dickinson. 2017b. Evolution of a split RNA polymerase as a versatile biosensor platform. Nat. Chem. Biol. 13:1–27. Nature Publishing Group.

Ramsey, G., and C. H. Pence. 2016. Chance in Evolution. The University of Chicago Press.

Reich, A., C. Dunn, K. Akasaka, and G. Wessel. 2015. Phylogenomic analyses of echinodermata support the sister groups of Asterozoa and Echinozoa. PLoS One 10:1–11.

Riesgo, A., N. Farrar, P. J. Windsor, G. Giribet, and S. P. Leys. 2014. The analysis of eight transcriptomes from all poriferan classes reveals surprising genetic complexity in sponges. Mol. Biol. Evol. 31:1102–1120.

Risso, V. A., F. Manssour-Triedo, A. Delgado-Delgado, R. Arco, A. Barroso-delJesus, A. Ingles-Prieto, R. Godoy-Ruiz, J. A. Gavira, E. A. Gaucher, B. Ibarra-Molero, and J. M. Sanchez-Ruiz. 2015. Mutational studies on resurrected ancestral proteins reveal conservation of site-specific amino acid preferences throughout evolutionary history. Mol. Biol. Evol. 32:440–455.

Sailer, Z. R., M. J. Harms, A. Dean, D. Usmanova, A. Mishin, and G. Sharonov. 2017. High-order epistasis shapes evolutionary trajectories. PLOS Comput. Biol. 13:e1005541.

Salverda, M. L. M., E. Dellus, F. a Gorter, A. J. M. Debets, J. van der Oost, R. F. Hoekstra, D. S. Tawfik, and J. A. G. M. de Visser. 2011. Initial mutations direct alternative pathways of protein evolution. PLoS Genet. 7:e1001321.

Schrödinger, L. 2018. The {PyMOL} Molecular Graphics System, Version∼2.0.7.

Shah, P., J. B. Plotkin, D. M. McCandlish, and J. B. Plotkin. 2015. Contingency and entrenchment in protein evolution under purifying selection. Proc. Natl. Acad. Sci. 112:E3226–E3235.

Shubin, N., C. Tabin, and S. Carroll. 2009. Deep homology and the origins of evolutionary novelty. Nature 457:818–823.

Smith, J. J., N. Timoshevskaya, C. Ye, C. Holt, M. C. Keinath, H. J. Parker, M. E. Cook, J. E. Hess, S. R. Narum, F. Lamanna, H. Kaessmann, V. A. Timoshevskiy, C. K. M. M. Waterbury, C. Saraceno, L. M. Wiedemann, S. M. C. C. Robb, C. Baker, E. E. Eichler, D. Hockman, T. Sauka-Spengler, M. Yandell, R. Krumlauf, G. Elgar, and C. T. Amemiya. 2018. The sea lamprey germline genome provides insights into programmed genome rearrangement and vertebrate evolution. Nat. Genet. 50:270–277. Springer US.

Somero, G. N. 1995. Proteins and Temperature. Annu. Rev. Physiol. 43–68.

Spor, A., D. J. Kvitek, T. Nidelet, J. Martin, J. Legrand, C. Dillmann, A. Bourgais, D. de Vienne, G. Sherlock, and D. Sicard. 2013. Phenotypic and Genotypic Convergences Are Influenced By Historical Contingency and Environment in Yeast. Evolution 1–54.

Starr, T. N., J. M. Flynn, P. Mishra, D. N. A. Bolon, and J. W. Thornton. 2018. Pervasive contingency and entrenchment in a billion years of Hsp90 evolution. Proc. Natl. Acad. Sci. 115:4453–4458.

Starr, T. N., L. K. Picton, and J. W. Thornton. 2017. Alternative evolutionary histories in the sequence space of an ancient protein. Nature 549:409–413. Nature Publishing Group.

Storz, J. F. 2016. Causes of molecular convergence and parallelism in protein evolution. Nat. Rev. Genet. 17:239–250. Nature Publishing Group.

Takechi, M., M. Takeuchi, K. G. Ota, O. Nishimura, M. Mochii, K. Itomi, N. Adachi, M. Takahashi, S. Fujimoto, H. Tarui, M. Okabe, S. Aizawa, and S. Kuratani. 2011. Overview of the transcriptome profiles identified in hagfish, shark, and bichir: Current issues arising from some nonmodel vertebrate taxa. J. Exp. Zool. Part B Mol. Dev. Evol. 316 B:526–546.

Thornton, J. W. 2004. Resurrecting ancient genes: experimental analysis of extinct molecules. Nat. Rev. Genet. 5:366–375.

Travisano, M., J. A. Mongold, A. F. Bennett, R. E. Lenski, M. Travisano, J. A. Mongold, and A. F. Bennett. 1995. Experimental Tests of the Roles of Adaptation, Chance, and History in Evolution. Science 267:87–90.

van Ditmarsch, D., K. E. Boyle, H. Sakhtah, J. E. Oyler, C. D. Nadell, É. Déziel, L. E. P. Dietrich, and J. B. Xavier. 2013. Convergent evolution of hyperswarming leads to impaired biofilm formation in pathogenic bacteria. Cell Rep. 4:697–708.

Waterhouse, A., M. Bertoni, S. Bienert, G. Studer, G. Tauriello, R. Gumienny, F. T. Heer, T. A. P. De Beer, C. Rempfer, L. Bordoli, R. Lepore, and T. Schwede. 2018. SWISS-MODEL: Homology modelling of protein structures and complexes. Nucleic Acids Res. 46:W296–W303. Oxford University Press.

Weir, B. S., and C. C. Cockerham. 1984. Estimating F-Statistics for the Analysis of Population Structure. Evolution 38:1358–1370.

Wichman, H. A., M. R. Badgett, L. A. Scott, C. M. Boulianne, and J. J. Bull. 1999. Different trajectories of parallel evolution during viral adaptation. Science 285:422–424.

Wu, N. C., A. J. Thompson, J. Xie, C. W. Lin, C. M. Nycholat, X. Zhu, R. A. Lerner, J. C. Paulson, and I. A. Wilson. 2018. A complex epistatic network limits the mutational reversibility in the influenza hemagglutinin receptor-binding site. Nat. Commun. 9. Springer US.

Wünsche, A., D. M. Dinh, R. S. Satterwhite, C. D. Arenas, D. M. Stoebel, and T. F. Cooper. 2017. Diminishing-returns epistasis decreases adaptability along an evolutionary trajectory. Nat. Ecol. Evol. 1:1–6.

Wyffels, J., B. L. King, J. Vincent, C. Chen, C. H. Wu, and S. W. Polson. 2014. SkateBase, an elasmobranch genome project and collection of molecular resources for chondrichthyan fishes. F1000Research 3.

Yokoyama, S., T. Tada, H. Zhang, and L. Britt. 2008. Elucidation of phenotypic adaptations: Molecular analyses of dim-light vision proteins in vertebrates. Proc. Natl. Acad. Sci. U. S. A. 105:13480–13485.

Závodszky, P., J. Kardos, Á. Svingor, and G. A. Petsko. 1998. Adjustment of conformational flexibility is a key event in the thermal adaptation of proteins. Proc. Natl. Acad. Sci. U. S. A. 95:7406–7411.

Zerbino, D. R., P. Achuthan, W. Akanni, M. R. Amode, D. Barrell, J. Bhai, K. Billis, C. Cummins, A. Gall, C. G. Girón, L. Gil, L. Gordon, L. Haggerty, E. Haskell, T. Hourlier, O. G. Izuogu, S. H. Janacek, T. Juettemann, J. K. To, M. R. Laird, I. Lavidas, Z. Liu, J. E. Loveland, T. Maurel, W. McLaren, B. Moore, J. Mudge, D. N. Murphy, V. Newman, M. Nuhn, D. Ogeh, C. K. Ong, A. Parker, M. Patricio, H. S. Riat, H. Schuilenburg, D. Sheppard, H. Sparrow, K. Taylor, A. Thormann, A. Vullo, B. Walts, A. Zadissa, A. Frankish, S. E. Hunt, M. Kostadima, N. Langridge, F. J. Martin, M. Muffato, E. Perry, M. Ruffier, D. M. Staines, S. J. Trevanion, B. L. Aken, F. Cunningham, A. Yates, and P. Flicek. 2018. Ensembl 2018. Nucleic Acids Res. 46:D754–D761.

Zhang, J., R. E. Campbell, A. Y. Ting, and R. Y. Tsien. 2002. Creating new fluorescent probes for cell biology. Nat. Rev. Mol. Cell Biol. 3:906–18.

Zheng, J., J. L. Payne, and A. Wagner. 2019. Cryptic genetic variation accelerates evolution by opening access to diverse adaptive peaks. Science 365:347–353.

Zhou, H., B. Sathyamoorthy, A. Stelling, Y. Xu, Y. Xue, Y. Z. Pigli, D. A. Case, P. A. Rice, and H. M. Al-Hashimi. 2019. Characterizing Watson-Crick versus Hoogsteen Base Pairing in a DNA-Protein Complex Using Nuclear Magnetic Resonance and Site-Specifically 13C- and 15N-Labeled DNA. Biochemistry 58:1963–1974.

Zhu, X., Y. Guan, A. Signore, C. Natarajan, S. DuBay, Y. Cheng, N. Han, G. Song, Y. Qu, H. Moriyama, F. Hoffmann, A. Fago, F. Lei, and J. Storz. 2018. Divergent and parallel routes of biochemical adaptation in high-altitude passerine birds from the Qinghai-Tibet Plateau. Proc. Natl. Acad. Sci. 115:1865–1870.

Zinkus-Boltz, J., C. Devalk, and B. C. Dickinson. 2019. A Phage-Assisted Continuous Selection Approach for Deep Mutational Scanning of Protein-Protein Interactions. ACS Chem. Biol. 14:2757–2767.

